# Virtual ChIP-seq: predicting transcription factor binding by learning from the transcriptome

**DOI:** 10.1101/168419

**Authors:** Mehran Karimzadeh, Michael M. Hoffman

## Abstract

**Motivation:** Identifying transcription factor binding sites is the first step in pinpointing non-coding mutations that disrupt the regulatory function of transcription factors and promote disease. ChIP-seq is the most common method for identifying binding sites, but performing it on patient samples is hampered by the amount of available biological material and the cost of the experiment. Existing methods for computational prediction of regulatory elements primarily predict binding in genomic regions with sequence similarity to known transcription factor sequence preferences. This has limited efficacy since most binding sites do not resemble known transcription factor sequence motifs, and many transcription factors are not even sequence-specific.

**Results:** We developed Virtual ChIP-seq, which predicts binding of individual transcription factors in new cell types using an artificial neural network that integrates ChIP-seq results from other cell types and chromatin accessibility data in the new cell type. Virtual ChIP-seq also uses learned associations between gene expression and transcription factor binding at specific genomic regions. This approach outperforms methods that predict TF binding solely based on sequence preference, predicting binding for 36 transcription factors (Matthews correlation coefficient > 0.3).

**Availability:** The datasets we used for training and validation are available at https://virchip.hoffmanlab.org. We have deposited in Zenodo the current version of our software (http://doi.org/10.5281/zenodo.1066928), datasets (http://doi.org/10.5281/zenodo.823297), predictions for 36 transcription factors on Roadmap Epigenomics cell types (http://doi.org/10.5281/zenodo.1455759), and predictions in Cistrome as well as ENCODE-DREAM *in vivo* TF Binding Site Prediction Challenge (http://doi.org/10.5281/zenodo.1209308).

## 1 Introduction

Transcription factor (TF) binding regulates gene expression. Each TF can harmonize expression of many genes by binding to genomic regions that regulate transcription. Cellular machinery utilizes these master regulators to regulate key cellular processes and adapt to environmental stimuli. Alteration in sequence or quantity of a given TF can impact expression of many genes. In fact, these alterations can be the primary cause of hereditary disorders, complex disease, autoimmune defects, and cancer^1^.

TFs bind to accessible chromatin based on weak non-covalent interactions between amino acid residues and nucleic acids. DNA’s primary structure (sequence)^2^, secondary structure (shape)^3^, and tertiary structure (conformation)^4^ all play roles in TF binding. Many TFs form a complex with others as well as chromatin-binding proteins and therefore bind to DNA indirectly. Some TFs also have different isoforms and undergo various post-translational modifications. *In vitro* assays, such as high throughput systematic evolution of ligands by exponential enrichment (HT-SELEX)^5^ and protein binding microarrays^6^, have provided a compelling understanding of context-independent TF sequence and shape preference^7^. Yet, for the aforementioned reasons, performance of models trained on these *in vitro* data are poor when applied on *in vivo* experiments^8,9^. To address this challenge, we must explore how to better model DNA shape, TF-TF interactions, and context-dependent TF binding.

Chromatin immunoprecipitation and sequencing (ChIP-seq)^10^ and similar methods, such as ChIP-exo^11^ and ChIP-nexus^12^, can map the presence of a given TF in the genome of a biological sample. To map TFs, these assays require a minimum of 1,000,000 to 100,000,000 cells, depending on properties of the TF itself and available antibodies. Such large numbers of cells are not often available from clinical samples. Therefore, it is impossible to systematically assess TF binding in most disease systems. Assessing chromatin accessibility through transposase-accessible chromatin using sequencing (ATAC-seq)^13^, however, requires only hundreds or thousands of cells. One can obtain this many cells from many more clinical samples. While chromatin accessibility does not determine TF binding, several methods use this information together with knowledge of TF sequence preference, genomic conservation, and other genomic features to predict TF binding ^14,15,16^.

Predicting TF binding with motif discovery tools within chromatin accessible regions has helped us understand the role of several TFs in various disease. For example, He et al.^17^ used motif discovery tools to identify the role of OCT1 and NKX3-1 after prolonged androgen stimulation in prostate cancer. Similarly, Bailey et al.^18^ discovered that a known breast cancer risk single nucleotide polymorphism (SNP) upstream of *ESR1* disrupts GATA3 binding and enhances expression of *ESR1*. We propose that using more accurate tools to predict TF binding will allow understanding the role of TF binding in more contexts.

Previous studies have used various approaches to predict TF binding. Several methods use unsupervised approaches such as hierarchical mixture models^14^ or hidden Markov models^15^ to identify transcription factor footprint using chromatin accessibility data. These approaches use sequence motif scores to attribute footprints to different transcription factors. Convolutional neural network models can boost precision by learning sequence preferences from *in vivo*, rather than *in vitro* data^20,21^,. Variation in sequence specificity and cooperative binding of some transcription factors prevents these methods from accurately predicting binding of all transcription factors. A more recent approach uses matrix completion to impute TF binding using a 3-mode tensor representing genomic positions, cell types, and TF binding^22^. This method doesn’t rely on sequence specificity, but can only predict TF binding in well-studied cell types with many ChIP-seq datasets. This means one cannot use it to predict binding in a cell type where ChIP-seq is not possible, such as limited clinical samples.

Identifying the best approach for predicting TF binding remains a challenge, because most studies use different benchmarking approaches. For example, one earlier study^14^ only assesses prediction on genomic regions that match the TF’s sequence motif. By excluding ChIP-seq peaks not matching the TF’s sequence motif from benchmarking, it underestimates false negative peaks and overestimates prediction accuracy. Most previous studies benchmark their predictions using the area under receiver operating characteristic curve (auROC) statistic^22,23,24^. When test data is imbalanced, meaning it has very different numbers of positive and negative examples, using auROC misleads evaluators^25,26^. Unfortunately, the TF binding status of genomic regions is highly imbalanced, making auROC alone a poor metric for evaluating TF binding prediction. Evaluation is further complicated by wildly varying prediction performance across different TFs. Recently, the ENCODE-DREAM *in vivo* TF Binding Site Prediction Challenge (DREAM Challenge) introduced guidelines for assessing TF binding prediction^27^. They recommend reporting both auROC, which assesses false negative predictions and the area under precision-recall curve (auPR), which also assesses false positives.

RNA-seq allows us to obtain transcriptome data from samples with small cell counts, including patient samples. We hypothesized that we could leverage the transcriptome to better predict TF binding. Previous methods have predicted gene expression using information on active regulatory elements^28,29,30^. Others have predicted chromatin accessibility using gene expression data^31^, or used differences in gene expression to identify statistically significant sequence motifs in specific conditions^32^, but they haven’t predicted *in vivo* TF binding using transcriptome data, as we do below.

Here, we introduce Virtual ChIP-seq, a novel method for more accurate prediction of TF binding. Virtual ChIP-seq predicts TF binding by learning from publicly available ChIP-seq experiments. Unlike Qin and Feng^23^, it can do this in new cell types with no existing ChIP-seq data. Virtual ChIP-seq also learns from other data such as genomic conservation, and the association of gene expression with TF binding.

Virtual ChIP-seq also accurately predicts the locations of DNA-binding proteins without known sequence preference. This would be impossible for most existing methods, which rely on sequence preference. Strictly speaking, only some of these proteins are TFs. As Lundberg et al. ^33^, we use the term *chromatin factors* in this paper to refer to factors subject to ChIP.

Virtual ChIP-seq predicted binding of 36 chromatin factors in new cell types with a minimum Matthews correlation coefficient (MCC) of 0.3. Eight of these chromatin factors (GTF2F1, HCFC1, HDAC2, NRF1, RAD21, SIN3A, SMC3, and TAF1) do not have DNA-binding domains and therefore are not TFs according to Lambert et al.^34^. These chromatin factors had minimum accuracy (fraction of all predictions that were correct) of 0.99 and minimum specificity (fraction of negative predictions that were correct) of 0.99. Precision (fraction of positive predictions that were correct) ranged between 0.16 and 0.78 (Table 1). We predicted binding of these 36 chromatin factors on 33 Roadmap Epigenomics^35^ cell types and provide these predictions as a track hub for community use (https://virchip.hoffmanlab.org).

**Table 1:** Performance of Virtual ChIP-seq for 36 chromatin factors on validation cell types. Each row displays median values ± standard deviation of several performance metrics for prediction of a chromatin factor across 4 chromosomes for each available validation cell type. MCC: Matthews correlation coefficient, auROC: area under receiver-operating characteristic curve, auPR: area under precision-recall, *N* : number of validation cell types for 36 chromatin factors with MCC > 0.3. We reported auROC and auPR across all the validation cell types across all posterior probability cutoffs. Black chromatin factors: we found the posterior probability cutoff which maximized MCC in H1-hESC, and then reported F_1_, accuracy, and MCC of the other validation cell types.

| chromatin factor | F<sub>1</sub> | Accuracy | MCC | auROC | auPR | N |
| --- | --- | --- | --- | --- | --- | --- |
| **ATF2** | 0.270±0.002 | 0.990±0.001 | 0.314±0.008 | 0.917±0.026 | 0.443±0.022 | 1 |
| **BHLHE40** | 0.334±0.021 | 0.997±0.000 | 0.356±0.010 | 0.974±0.002 | 0.382±0.01 | 1 |
| **CEBPB** | 0.510±0.091 | 0.992±0.002 | 0.515±0.072 | 0.964±0.017 | 0.534±0.073 | 3 |
| **CHD2** | 0.399±0.038 | 0.998±0.000 | 0.406±0.034 | 0.950±0.012 | 0.386±0.046 | 1 |
| **CREB1** | 0.362±0.131 | 0.997±0.002 | 0.371±0.121 | 0.868±0.135 | 0.335±0.174 | 2 |
| **CTCF** | 0.667±0.126 | 0.995±0.004 | 0.675±0.092 | 0.985±0.050 | 0.841±0.108 | 6 |
| **E2F1** | 0.256±0.097 | 0.998±0.002 | 0.314±0.078 | 0.978±0.019 | 0.291±0.105 | 2 |
| **ELF1** | 0.431±0.047 | 0.997±0.001 | 0.456±0.038 | 0.949±0.042 | 0.493±0.066 | 2 |
| **ELK1** | 0.430±0.069 | 1.000±0.000 | 0.465±0.054 | 0.991±0.009 | 0.420±0.054 | 2 |
| **ESR1** | 0.372±0.103 | 0.993±0.006 | 0.430±0.049 | 0.883±0.033 | 0.461±0.019 | 2 |
| **FOS** | 0.333±0.027 | 0.997±0.001 | 0.393±0.020 | 0.861±0.004 | 0.394±0.008 | 1 |
| **FOSL1** | 0.319±0.006 | 0.994±0.001 | 0.316±0.006 | 0.929±0.006 | 0.272±0.012 | 1 |
| **FOXA1** | 0.433±0.082 | 0.997±0.004 | 0.492±0.072 | 0.981±0.022 | 0.568±0.117 | 3 |
| **GABPA** | 0.298±0.049 | 0.994±0.002 | 0.393±0.036 | 0.986±0.012 | 0.496±0.036 | 3 |
| **GTF2F1** | 0.235±0.120 | 0.996±0.001 | 0.312±0.070 | 0.985±0.015 | 0.191±0.081 | 2 |
| **HCFC1** | 0.459±0.021 | 0.999±0.000 | 0.487±0.024 | 0.990±0.005 | 0.515±0.044 | 2 |
| **HDAC2** | 0.303±0.033 | 0.986±0.005 | 0.370±0.018 | 0.948±0.051 | 0.281±0.040 | 2 |
| **HSF1** | 0.350±0.149 | 1.000±0.000 | 0.378±0.145 | 0.999±0.012 | 0.309±0.240 | 1 |
| **JUN** | 0.218±0.127 | 0.998±0.001 | 0.311±0.153 | 0.983±0.009 | 0.456±0.257 | 2 |
| **JUND** | 0.341±0.163 | 0.993±0.002 | 0.386±0.135 | 0.979±0.019 | 0.326±0.161 | 4 |
| **MAFK** | 0.354±0.041 | 0.997±0.001 | 0.423±0.028 | 0.989±0.005 | 0.513±0.103 | 3 |
| **MAX** | 0.400±0.045 | 0.996±0.002 | 0.444±0.059 | 0.961±0.012 | 0.491±0.111 | 3 |
| **MAZ** | 0.370±0.025 | 0.997±0.001 | 0.422±0.019 | 0.987±0.005 | 0.493±0.070 | 2 |
| **MXI1** | 0.394±0.018 | 0.999±0.000 | 0.402±0.017 | 0.993±0.004 | 0.381±0.025 | 1 |
| **NRF1** | 0.658±0.042 | 1.000±0.000 | 0.664±0.038 | 0.994±0.014 | 0.720±0.051 | 3 |
| **RAD21** | 0.593±0.062 | 0.996±0.002 | 0.626±0.056 | 0.983±0.033 | 0.740±0.095 | 3 |
| **REST** | 0.482±0.120 | 0.999±0.001 | 0.493±0.091 | 0.985±0.008 | 0.567±0.095 | 3 |
| **SIN3A** | 0.389±0.048 | 0.998±0.002 | 0.394±0.029 | 0.966±0.004 | 0.411±0.037 | 3 |
| **SMC3** | 0.733±0.016 | 0.999±0.000 | 0.734±0.016 | 0.998±0.001 | 0.792±0.018 | 1 |
| **SRF** | 0.353±0.060 | 0.998±0.001 | 0.364±0.070 | 0.982±0.008 | 0.365±0.115 | 2 |
| **TAF1** | 0.378±0.073 | 0.999±0.001 | 0.437±0.097 | 0.987±0.009 | 0.490±0.168 | 3 |
| **TEAD4** | 0.344±0.061 | 0.990±0.002 | 0.385±0.020 | 0.967±0.023 | 0.343±0.019 | 2 |
| **TP53** | 0.275±0.103 | 1.000±0.000 | 0.382±0.086 | 1.000±0.008 | 0.660±0.222 | 1 |
| **USF1** | 0.353±0.047 | 0.993±0.001 | 0.382±0.040 | 0.891±0.012 | 0.372±0.046 | 1 |
| **USF2** | 0.410±0.040 | 0.999±0.000 | 0.427±0.028 | 0.982±0.007 | 0.437±0.032 | 1 |
| **YY1** | 0.397±0.049 | 0.996±0.001 | 0.408±0.058 | 0.945±0.043 | 0.417±0.104 | 2 |

## 2 Results

### 2.1 Sequence motifs are absent in most TF binding sites

#### 2.1.1 Most ChIP-seq peaks lack the TF’s relevant sequence motif

Many computational tools predict TF binding using sequence preference data^14,15^. Most tools represent TF sequence preference in position weight matrix (PWM) format. PWMs encode the likelihood for presence of each nucleotide at different positions of a sequence motif. With tools such as FIMO^36^, we can efficiently search and rank genomic regions that match TF sequence motifs.

One cannot determine a TF’s binding sites based solely on its sequence preference. We can identify some additional properties, such as co-binding partners, from high-throughput experiments. For other properties, such as post-translational modifications to the TF, we lack corresponding large-scale data. Many post-translational modifications affect cellular localization, binding partners, and DNA-recognition of chromatin factors^37^. Therefore, we expect existing computational prediction methods to be more accurate for chromatin factors where post-translational modifications and cobinding partners contribute less to TF binding. For chromatin factors with more complex biology, however, we expect computational prediction methods to fail.

Using ChIP-seq data on 201 chromatin factors in 54 different cell types, we investigated whether the majority of binding sites matched the sequence motif of the same TF. Among these 201 proteins, 76 lacked a sequence motif in JASPAR (Figure 1a, Supplementary Table 1). Some of these motif-free proteins, such as EZH2 and HDAC, are chromatin-binding proteins rather than true TFs. For simplicity in describing the prediction task, we refer to them as chromatin factors. Others are TFs without known sequence preference. For sequence-specific TFs, the fraction of peaks that match a sequence motif ranges from 4.55% (for SIX5) to 94.2% (for CTCF) with a mean of 49.4% (Figure 1b).

**Figure 1:**
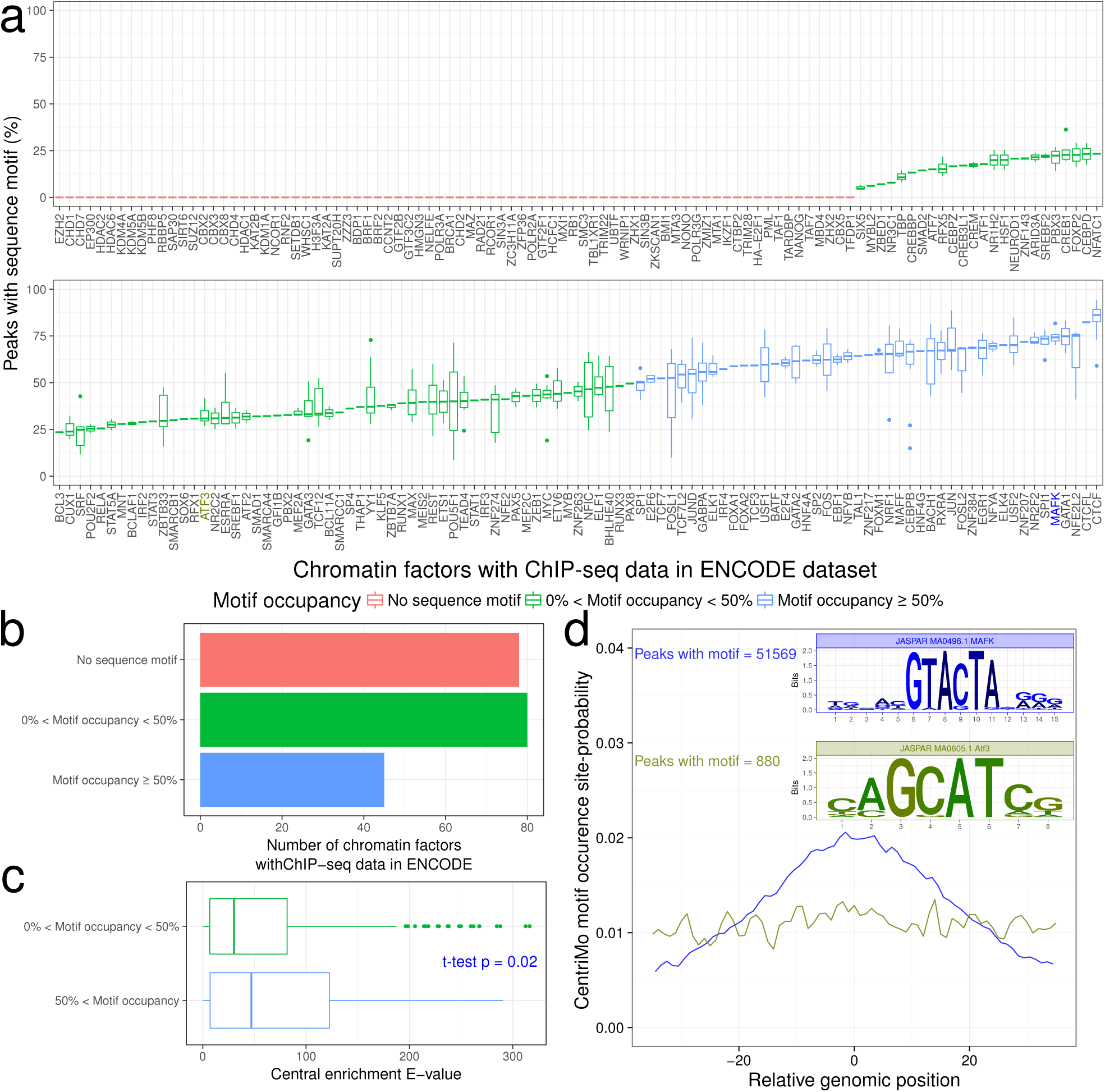
Most ChIP-seq peaks lack the TF’s sequence motif. **(a)** Fraction of ENCODE ChIP-seq peaks for a TF with any JASPAR sequence motif from the TF’s family. Boxplots show the distribution among datasets from different cell types and replicates. Horizontal line of boxplot: median. Box range: interquartile range (IQR). Whisker: most extreme value within quartile ± 1.5 IQR. Individual points: outliers beyond a whisker. **(b)** Number of factors without a sequence motif in JASPAR (red), TFs where less than 50% of peaks have the sequence motif (low motif occupancy, green), and TFs where 50% or more of peaks have the sequence motif (high motif occupancy, blue). **(c)** Central enrichment^19^ of a TF’s motif is lower for TFs with motif occupancy of less than 50% compared to TFs with motif occupancy of 50% or more. **(d)** For TFs with a small number of peaks matching sequence motif of the same TF, such as ATF3, central enrichment of the motif is also low. In contrast, most MAFK peaks both contain its sequence motif and show central enrichment.

#### 2.1.2 Many sequence motifs are not centrally enriched

Central enrichment measures how close a sequence motif occurs to a set of ChIP-seq peak summits. According to Bailey and Machanick^19^, high central enrichment indicates direct TF binding. We used CentriMo^19^ to measure central enrichment. We compared central enrichment between TFs with low motif occupancy (< 50% of ChIP-seq peaks contain the motif) and high motif occupancy (≥ 50% of peaks contain the motif; Figure 1c). TFs with low motif occupancy had weaker central enrichment (t-test; *p* = 0.02). For example, 30.87% of ATF3 peaks overlapped with the MA0605.1 JASPAR motif. ATF3 peaks also had lower central enrichment than MAFK peaks, which had 74.29% overlap with the MA0496.1 JASPAR motif (Figure 1d).

### 2.2 Model, performance, and benchmarking

#### 2.2.1 Datasets

Virtual ChIP-seq learns from the association of gene expression and chromatin factor binding in publicly available datasets. Our method requires ChIP-seq data of each chromatin factor in as many cell types as possible, with matched RNA-seq data from the same cell types. We used ChIP-seq data (from Cistrome DB^38^ and ENCODE^39^) and RNA-seq data (from CCLE^40^ and ENCODE^41^) to assess Virtual ChIP-seq’s binding predictions for 63 DNA-binding proteins in new cell types. We considered a 200 bp genomic bin as *bound* if it overlapped a peak summit with FDR < 10^−4^ (Methods).

In addition to benchmarking on our own held-out test cell types, we wanted to compare against the DREAM Challenge^27^. To do this, we also used their datasets, which include ChIP-seq data for 31 chromatin factors. For most of these chromatin factors, the DREAM Challenge held out test chromosomes instead of test cell types. The DREAM Challenge included ChIP-seq data for only 12 chromatin factors in completely held-out cell types. Completely holding out cell types better fits the real-world scenarios that require binding site prediction. Using the datasets we generated, we had matched data in enough cell types to train and validate models for 9 of these 12 chromatin factors (CTCF, E2F1, EGR1, FOXA1, GABPA, JUND, MAX, REST, and TAF1).

#### 2.2.2 Learning from the transcriptome

Different cell types have distinct transcriptomic and epigenomic states^42^. Changing gene expression levels can affect patterns of chromatin factor binding and chromatin structure. We hypothesized that some gene expression changes would lead to consistent and observable changes in chromatin factor binding. As an extreme example, eliminating expression of a chromatin factor would eventually eliminate binding of that chromatin factor genome-wide. Other changes in gene expression could lead to competitive, cooperative, allosteric, and other indirect effects that would affect chromatin factor binding. To account for both direct and indirect effects of the expression of regulatory genes, one must model the dependency of each chromatin factor binding site on expression of all genes^31^. To exploit this model, we identified genes with significant positive or negative correlation with chromatin factor binding at any given genomic bin. We did this for genes all over the genome, irrespective of distance from the binding site.

For each chromatin factor, we created an *association matrix* measuring correlation between gene expression and binding of that chromatin factor in previously collected datasets (Figure 2a– c). In this matrix, each value corresponds to the Pearson correlation between ChIP-seq binding of that chromatin factor at one genomic bin and the expression level of one gene. We used missing values when there was no significant association between gene expression and chromatin factor binding (*p* > 0.1).

**Figure 2:**
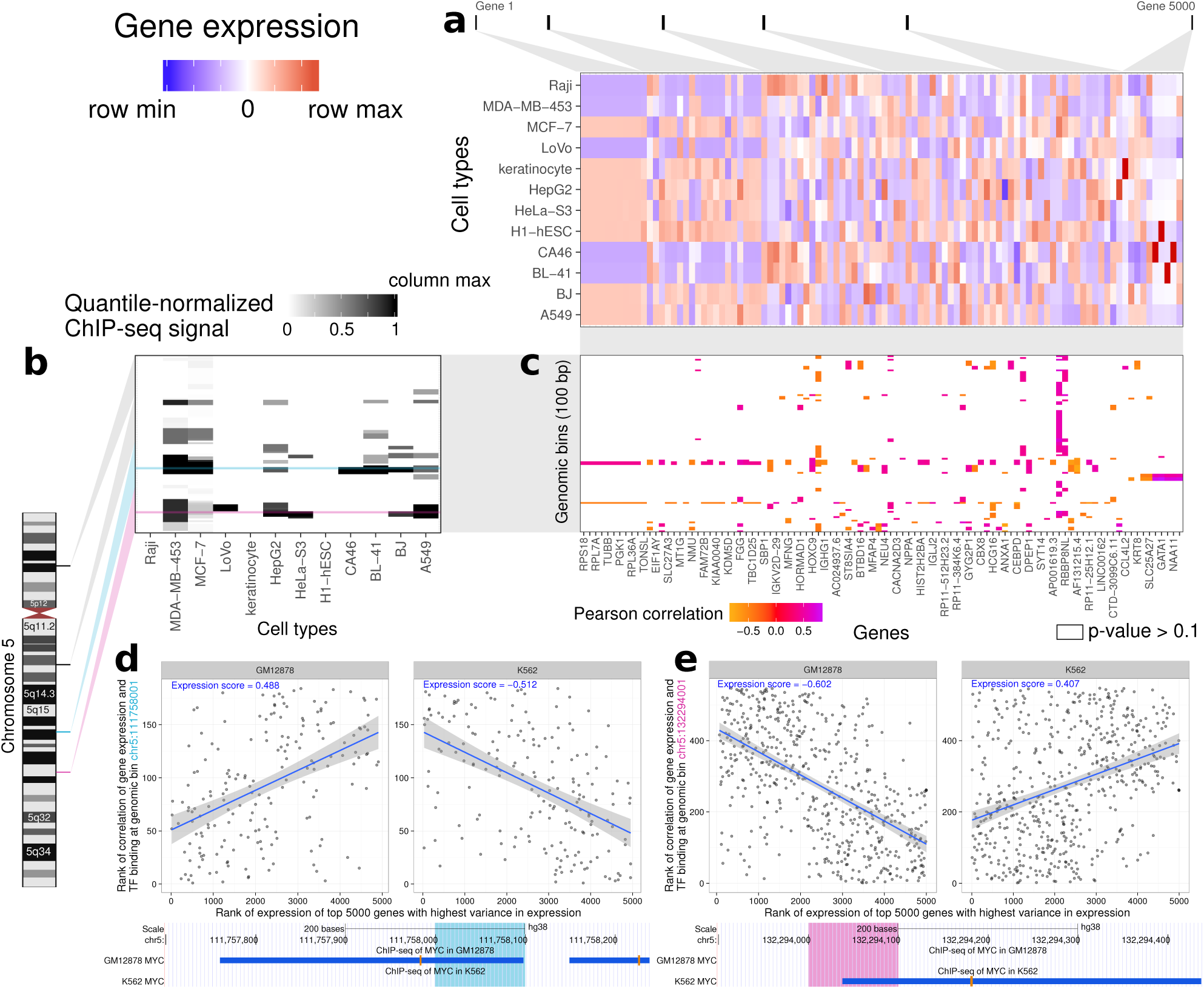
Virtual ChIP-seq learns from association of gene expression and chromatin factor binding at each genomic bin. This example shows Virtual ChIP-seq analysis for the MYC TF. **(a)** Gene expression levels for 5000 genes in 12 cell types. For simplicity of visualization, we showed only 100 of these genes in the matrix and labeled only every other gene. We ranked RNA-seq RPKM expression values within each cell type. This matrix shows a subset of 5000 high-variance genes, sorted by variance of each gene’s expression between cell types. Blue: row minimum; white: median expression; red: row maximum. **(b)** ChIP-seq signal for 100 bp bins in 12 cell types, taken from four larger regions (25 bins each) on chromosome 5. We quantile-normalized ChIP signal from MACS software among cell types. This matrix shows a subset of the 54,037 bins on chromosome 5 which have TF binding in at least one training cell type. White: column minimum (0.0); black: column maximum (1.0). Cyan: a region in the *NREP* locus with *MYC* binding in GM12878; magenta: a region upstream of *SLC22A4* with *MYC* binding in K562. **(c)** Association matrix: gene expression–ChIP signal correlation between 100 genomic bins and 5000 high-variance genes. This is a subset of the larger 54,037 × 5,000 association matrix for chromosome 5. Each cell shows the Pearson correlation for 12 cell types between expression for a particular gene and ChIP signal at a particular genomic bin. Orange: negative correlation; white: p-value of Pearson correlation greater than 0.1 (NA); Purple: positive correlation. **(d)** (*Top*) Expression score plots for a 100 bp bin in the *NREP* locus. Each plot has one point for each of 184 genes with non-NA correlation values at that bin in the association matrix. Each point displays the rank of correlation value for that gene among one row of the association matrix against the rank of expression for that gene among 5000 high-variance genes in (*left*) GM12878 and (*right*) K562 cell types. The expression score at a bin for a cell type is Spearman’s rank correlation coefficient *ρ* between those two ranks. Blue line: best linear fit to data; grey region: 95% confidence interval of the fit. (*Bottom*) UCSC Genome Browser display of 550 bp around that region. Blue rectangle: *MYC* ChIP-seq peak in GM12878 or K562. Here, *MYC* binds only in GM12878. **(e)** Expression score plot and Genome Browser display for a 100 bp bin upstream of *SLC22A4*. Here, *MYC* binds only in K562.

Power analysis (Methods) identified which correlations the *p* > 0.1 cutoff would exclude depending on the number of available cell types with matched ChIP-seq and RNA-seq data. For CTCF, which had the largest number of cell types available—21 cell types with matched ChIP-seq and RNA-seq—this cutoff provided 80% power to detect an absolute value of Pearson correlation *|r|* ≥ 0.52. Many chromatin factors had only 5 cell types with matched data and the cutoff provided 80% power to detect only larger correlations, *|r|* ≥ 0.92.

We calculated an *expression score* for a chromatin factor in a new cell type using the association matrix and RNA-seq data for the new cell type, but no ChIP-seq data. The expression score is the Spearman correlation between the non-NA values for that genomic bin in the association matrix and the expression levels of those genes in the new cell type (Figure 2d, Figure 3a). We used the rank-based Spearman correlation to make the score robust against slight differences in analytical methodology used to estimate gene expression. We used the expression score as one of the variables in a multi-layer perceptron (Methods). Compared to using the high-dimensional and sparse association matrix as an input to the multi-layer perceptron, this approach has several advantages. Most importantly, the expression score is dependent on the transcriptome of the new cell type. Other advantages include a much lower number of parameters to optimize, and avoiding the problem of NA values in the association matrix.

**Figure 3:**
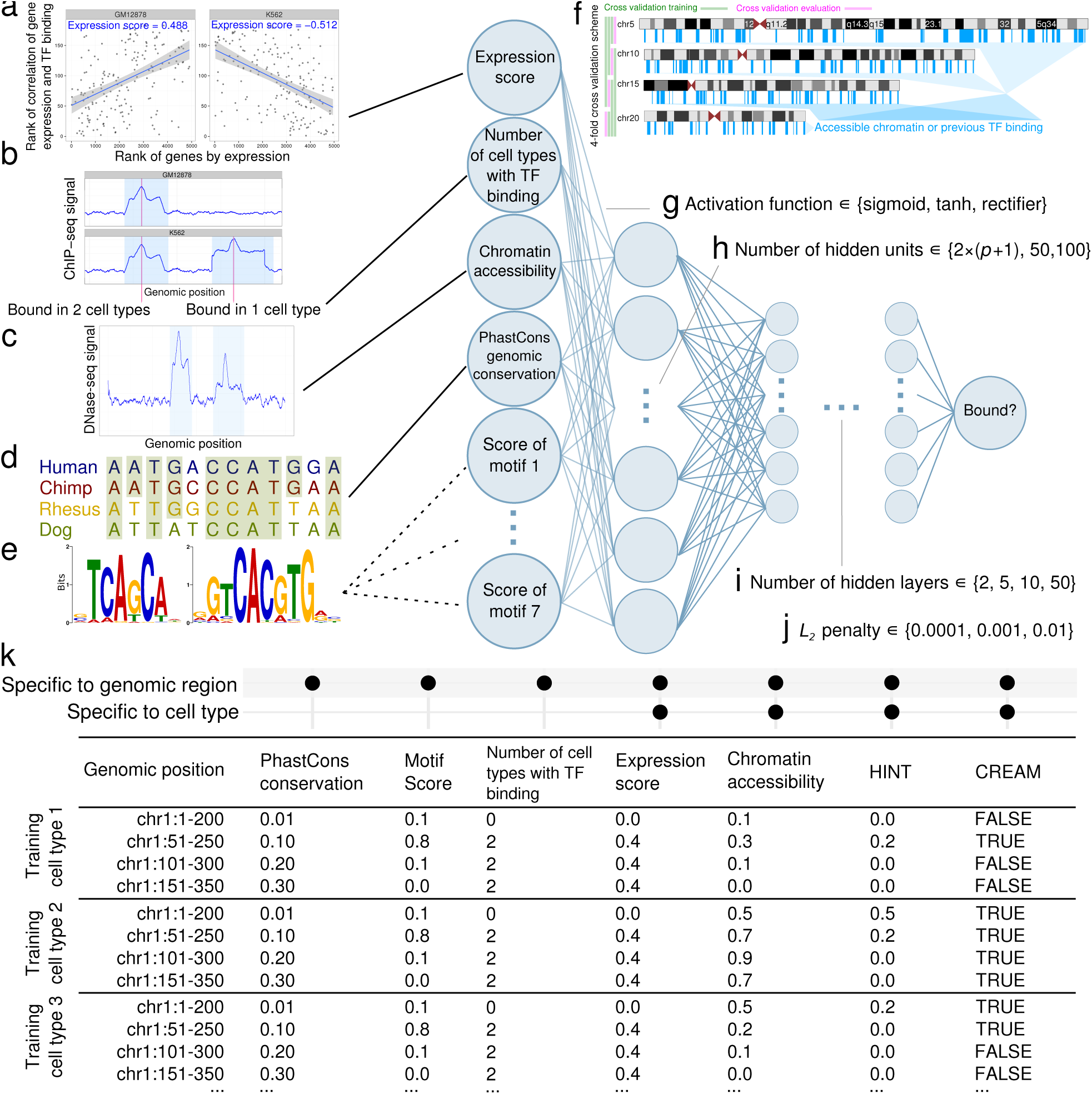
Optimizing and training a multi-layer perceptron. We used a number of features to predict chromatin factor binding in each bin. These include **(a)** expression score (Figure 2d–e), **(b)** the number of training cell types with binding of that chromatin factor, **(c)** chromatin accessibility, **(d)** PhastCons genomic conservation in placental mammals, and **(e)** any sequence motif corresponding to that TF in the JASPAR database. In JASPAR, some chromatin factors have no sequence motifs, while others have up to seven different sequence motifs. This led to a number of features *p ∈* [4, 11], excluding features from HINT footprints or CREAM peaks not used in the main model. **(f)** For each chromatin factor, we trained a multi-layer perceptron using these features for selected bins in four chromosomes (5, 10, 15, and 20). Specifically, we selected bins with accessible chromatin or ChIP-seq signal in at least one training cell type (selected regions with vertical blue bars are for illustration purpose). To optimize hyperparameters, we repeated the training process with different hyperparameters using four-fold cross validation, excluding one chromosome at a time. For each chromatin factor, we performed a grid search over **(g)** activation function (sigmoid, tanh, and rectifier), **(h)** number of hidden units per layer (2(*p* +1), 50, or 100), **(i)** number of hidden layers (2, 5, 10, or 50), and **(j)** *L*_2_ regularization penalty (0.0001, 0.001, or 0.01). We chose the quadruple of hyperparameters which resulted in the highest mean Matthews correlation coefficient (MCC) over all four chromosomes. **(k)** Schematic of the matrix of input features for training the multi-layer perceptron. We used the input features from all the available training cell types to train the multi-layer perceptron.

#### 2.2.3 Learning from other predictive features

We included a number of other predictive features beyond expression score. Virtual ChIP-seq includes as input for each genomic bin the frequency of the chromatin factor’s presence in existing ChIP-seq data (Figure 3b). Since most chromatin factor binding occurs within accessible chromatin^43^, we also used evidence of chromatin accessibility from DNase-seq or ATAC-seq (Figure 3c).

While many intra-species genomic differences lie in the non-coding genome^44^, we expect some regulatory elements to be conserved among closely related species. Previous studies highlight the association of genomic conservation and chromatin factor binding in organisms as simple as yeast^45^ or as complex as human^46^. To learn from patterns of genomic conservation, we used PhastCons^47,48^ scores from a 7-way primate and placental mammal comparison (http://hgdownload.cse.ucsc.edu/goldenPath/hg38/phastCons7way) in our model (Figure 3d).

We used sequence motif score where available (Figure 3e; see Methods). Relying only on TF sequence preference, however, would prevent accurate prediction of most true TF binding sites^9^ (Figure 1). For each TF, we represented sequence preference using the FIMO score of JASPAR sequence motifs of that TF or a similar TF. JASPAR has no motif for some chromatin factors, such as EP300. Where JASPAR has more than one motif for a TF, additional motifs often represent different versions of the motif such as SREBF2 (MA0596.1) and SREBF2-var2 (MA0828.1). In some cases, the additional motif represents a preference of a cooperative TF heterodimer, such as MAX-MYC (MA0059.1). Regardless of reason, we included all of each TF’s motifs as features in its model (Supplementary Table 2).

We also investigated potential improvements by adding a couple of additional integrative features available for a limited number of chromatin factors and cell types (Supplementary Table 2). First, we used the output of Hidden Markov model-based Identification of TF footprints (HINT)^15^ which identifies TF footprints within accessible chromatin. Second, we used a boolean feature indicating overlap of each genomic bin with clusters of chromatin accessibility peaks identified by CREAM^49^.

#### 2.2.4 Selecting hyperparameters and training

We created an input matrix with rows corresponding to 200 bp genomic windows and columns representing the features described above. Specifically, these features included expression score (Figure 3a), previous evidence of binding of chromatin factor of interest in publicly available ChIP-seq data (Figure 3b), chromatin accessibility (Figure 3c), genomic conservation (Figure 3d), sequence motif scores (Figure 3e), HINT footprints, and CREAM peaks. We used sliding genomic bins with 50 bp shifts, where most 200 bp bins overlap six other bins. This provided a maximum resolution of 50 bp in binding prediction. This resulted in a sparse matrix with 60,620,768 rows representing each bin in the GRCh38 genome assembly^50^. The sparse matrix used in the main model had between 4 and 11 columns, depending on the number of available sequence motifs. When we added HINT footprints and CREAM peaks, the matrix had between 6 and 13 columns instead. We trained on an imbalanced subset of genomic regions which had chromatin factor binding or chromatin accessibility (FDR < 10^−4^) in any of the training cell types. To speed the process of training and evaluation, we further limited training input data to four chromosomes (chr5, chr10, chr15, and chr20). For validation, however, we used data from these same four chromosomes in completely different cell types held out from training. We evaluated the performance on all of the 9,635,407 bins in these four chromosomes (Figure 3f), not just those with prior evidence of chromatin factor binding or chromatin accessibility.

To build a generalizable classifier that performs well on new cell types with only transcriptome and chromatin accessibility data, we concatenated input matrices from 12 training cell types: A549, GM12878, HepG2, HeLa-S3, HCT-116, BJ, Jurkat, NHEK, Raji, Ishikawa, LNCaP, and T47D (Supplementary Table 3).

#### 2.2.5 The multi-layer perceptron

The multi-layer perceptron is a fully connected feed-forward artificial neural network^51^. Our multi-layer perceptron assumes binding at each genomic window is independent of upstream and down-stream windows (Figure 3). For each chromatin factor, we trained the multi-layer perceptron with adaptive momentum stochastic gradient descent^52^ and a minibatch size of 200 samples. We used 4-fold cross validation to optimize hyperparameters including activation function (Figure 3g), number of hidden units per layer (Figure 3h), number of hidden layers (Figure 3i), and *L*_2_ regularization penalty (Figure 3j). For training, we only used genomic bins which overlapped chromatin accessibility peaks or previous evidence of chromatin factor binding in any of the training cell types. For assessing performance, however, we used all genomic bins of the chromosome (Methods). In each cross validation fold, we iteratively trained on 3 of the 4 chromosomes (5, 10, 15, and 20) at a time, and assessed performance in the remaining chromosome. We selected the model with the highest average Matthews correlation coefficient (MCC)^53^ after 4-fold cross validation. MCC incorporates all four categories of a confusion matrix and assesses performance well even on imbalanced datasets^54^. For 23 chromatin factors, the optimal model had 10 hidden layers. For another set of 23 chromatin factors, the optimal model had 5 hidden layers. For the final 17 chromatin factors, the optimal model had only 2 hidden layers.

For 57 out of the 63 chromatin factors examined, the best-performing model had 100 hidden units in each layer—the maximum number of hidden units per layer examined in the grid search. For the remaining 6 chromatin factors, the optimal model had 10–24 hidden units in each layer. Different activation functions—hyperbolic tangent (tanh) or rectifier—proved optimal for different chromatin factors (Supplementary Table 4).

We investigated if chromatin factors with the same DNA binding domain (as reported in Lambert et al.^34^) also have similar optimized hyperparameters. All C2H2 zinc finger TFs (EGR1, CTCF, MAZ, REST, YY1, ZBTB33, ZNF143, and ZNF274) had a rectifier activation function, 100 hidden units, and *L*_2_ regularization penalty of 10^−4^. The number of hidden layers ranged from 2 to 10. The other DNA binding domains which had more than 4 TFs in our datasets, bHLH and bZIP, did not have the same hyperparameter among their TFs (Supplementary Table 4). There was also no significant correlation between number of hidden layers, hidden units, or activation function with performance of the model in validation cell types.

#### 2.2.6 Virtual ChIP-seq predicts chromatin factor binding with high accuracy

We evaluated the performance of Virtual ChIP-seq in validation cell types (K562, PANC-1, MCF-7, IMR-90, H1-hESC, and primary liver cells) which we did not use in calculating the expression score, training the multi-layer perceptron, or optimizing hyperparameters. Before predicting in new cell types, we chose a posterior probability cutoff for use in point metrics such as accuracy and F_1_ score. When a chromatin factor had ChIP-seq data in more than one of the validation cell types, we chose the cutoff that maximizes MCC of that chromatin factor in H1-hESC cells. Then, we excluded H1-hESC when reporting threshold-requiring metrics. For these chromatin factors, we pre-set a posterior probability cutoff of 0.4, the mode of the cutoffs for other chromatin factors (Supplementary Table 5).

Virtual ChIP-seq predicts binding of 36 chromatin factors in validation cell types with MCC *>* 0.3, auROC *>* 0.9, and 0.3 < auPR < 0.8 (Figure 4a, Table 1, Supplementary Table 6).

**Figure 4:**
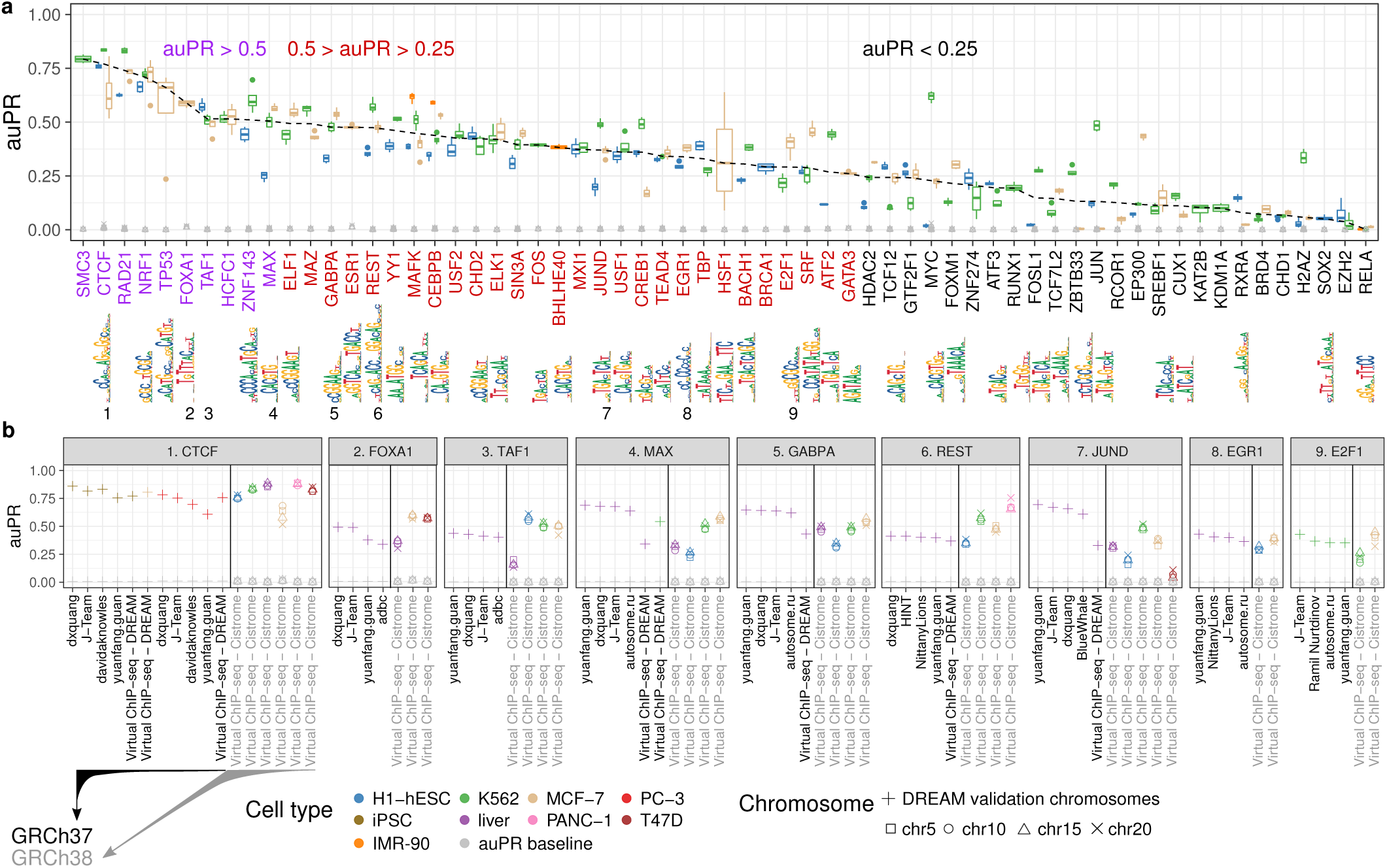
Virtual ChIP-seq predicts chromatin factor binding with high accuracy. Using ChIP-seq and RNA-seq data, we learned from the association of gene expression and chromatin factor binding for 63 chromatin factors. **(a)** Box plots show distribution of auPR among 4 chro-mosomes (5, 10, 15, and 20) for 63 chromatin factors assessed in four cell types (blue: H1-hESC; orange: IMR-90; green: K562; brown: MCF-7). Dashed line: medians. Grey shapes: prevalence of bound bins in the chromosome, the auPR baseline. Axis label colors categorize median auPR (purple: greater than 0.5, red: between 0.25 and 0.5, black: below 0.25). Sequence logos indicate one of a TF’s JASPAR motifs, when available. When multiple motifs existed, we displayed the shortest motif here. Numbers 1–9: The nine chromatin factors that the DREAM Challenge also evaluated in its final round. **(b)** We compared Virtual ChIP-seq’s performance to that of the top 4 performing methods in the DREAM Challenge across-cell type final round. For CTCF, MAX, GABPA, REST, and JUND, we had enough cell types to train and validate the performance of Virtual ChIP-seq on DREAM data. For these chromatin factors we trained on chromosomes 5, 10, 15, and 20 in training cell types and validated performance on merged data of chromosomes 1, 8, and 21 in validation cell types. For other chromatin factors, we trained the model and validated our performance using publicly available Cistrome and ENCODE data. auPR values are only directly comparable for the same cell type and test set. The black vertical line in each panel separates test sets based on genome assembly and source. Axis label color: reference genome assembly (black: GRCh37, grey: GRCh38).

#### 2.2.7 Virtual ChIP-seq correctly predicts binding sites in genomic locations not found in training data

We evaluated the performance of Virtual ChIP-seq for 63 chromatin factors with binding in validation cell types. For 59 of these chromatin factors, Virtual ChIP-seq predicted true chromatin factor binding in regions without conservation among placental mammals. For 44 out of 63 chromatin factors, Virtual ChIP-seq predicted true chromatin factor binding in regions without chromatin factor binding in any of the training ChIP-seq data. From these 63 chromatin factors, 43 are sequence-specific, and for all of these chromatin factors, Virtual ChIP-seq predicted true binding for regions that did not match the TF’s sequence motif. For 47 chromatin factors, Virtual ChIP-seq even correctly predicted chromatin factor binding in regions that didn’t overlap chromatin accessibility peaks (Supplementary Table 7). Most of these regions were frequently bound to the chromatin factor in publicly available ChIP-seq data. These predictions showed that the multi-layer perceptron learned to leverage multiple kinds of information and predict chromatin factor binding accurately, even in the absence of features required by previous generations of binding site classifiers.

#### 2.2.8 Comparison with DREAM Challenge

DREAM Challenge rules forbid using genomic conservation or ChIP-seq data as training features. This also excludes the expression score, as creating its association matrix relies on ChIP-seq data. The challenge also required training and validation on its own provided datasets. These datasets have ChIP-seq data in only a few cell types. This restricts Virtual ChIP-seq’s approach which leverages all publicly available datasets. The DREAM Challenge ChIP-seq datasets use only two replicates for each experiment and requires that peaks have a irreproducibility discovery rate (IDR)^55^ of less than 5%.

In these cases, we included peaks that pass a false discovery rate (FDR) threshold of 10^−4^ in at least two replicates (Supplementary Table 8).

The DREAM Challenge assessed participant entries by measuring performance on three validation chromosomes (chr1, chr8, and chr21), combined. To assess performance of Virtual ChIP-seq on DREAM Challenge data, we did the same. To assess performance on Cistrome data, however, we measured performance on each chromosome independently. This allowed us to examine the variance in performance among these chromosomes.

Although Virtual ChIP-seq used features not allowed in the DREAM Challenge, comparing with DREAM Challenge participants is the only sound way to show how any method including these features compares to the state of the art. Before the DREAM Challenge, TF binding prediction methods mostly reported performance measurements only in those parts of a chromosome where a method had more likelihood of success. The DREAM Challenge, like Virtual ChIP-seq, instead reports performance on the intended deployment domain of such methods: whole chromosomes. Leading DREAM Challenge methods potentially could improve their performance by including the features used by Virtual ChIP-seq. We compared Virtual ChIP-seq with DREAM Challenge results when we trained and validated on either Cistrome DB data or DREAM Challenge data.

#### 2.2.9 Prediction accuracy varies by transcription factor

The DREAM Challenge evaluates predictions on binding of 31 chromatin factors. The final submission round evaluates predictions for 12 chromatin factors in held-out cell types. The datasets we used, however, allow us to predict binding of 63 chromatin factors in new cell types. Of these chromatin factors, 41 are unique to our dataset and do not overlap any of the DREAM Challenge chromatin factors (Supplementary Table 9). The DREAM Challenge has data on the other 22 chromatin factors, but the challenge evaluated only 9 of these chromatin factors in its final round.

For CTCF, FOXA1, TAF1, and REST, Virtual ChIP-seq had a higher auPR in at least one validation cell type than any DREAM Challenge participant^56,57^. For EGR1 and E2F1, Virtual ChIP-seq performed better than at least one of the four top-performing methods of the challenge in one of the validation cell types (Figure 4b). DREAM Challenge and Cistrome ChIP-seq peak calls had different class imbalances, making auPR statistics not directly comparable (Supplementary Table 10). These imbalances were not always in the same direction. In FOXA1 peak calls in liver, for example, Cistrome called 0.12% of genomic bins bound to a chromatin factor, half the fraction of the DREAM Challenge (0.25%). Our predictions for FOXA1 binding in T47D and MCF-7 using Cistrome had a higher auPR than participants of DREAM Challenge for liver. The FOXA1 peak calls for these cell types also had a higher fraction of chromatin factor-bound genomic bins: 1.36% for MCF-7, and 0.39% for T47D. This opposed the smaller fraction of bins bound in Cistrome data in CTCF (in PANC-1, liver, and T47D), TAF1 (in liver, H1-hESC, K562, and T47D), and REST (in H1-hESC, K562, and PANC-1). The differences in class prevalence are both minor and in diverging directions. Because of this, they do not bias the baseline auPR of evaluation on Cistrome datasets in a particular direction when compared to evaluation on DREAM Challenge datasets.

The power of Virtual ChIP-seq to learn from the transcriptome data diminishes when fewer cell types are available, as in the DREAM Challenge data. Nonetheless, when trained on DREAM Challenge data, Virtual ChIP-seq outperformed 13/14 DREAM Challenge participants when predicting CTCF binding in PC-3 cells. When predicting CTCF binding in iPSC cells, Virtual ChIP-seq had a higher auPR than 8/14 Challenge participants. The Virtual ChIP-seq auPR for binding of REST in liver was also higher than that of 9/14 DREAM Challenge participants (Supplementary Table 11).

Virtual ChIP-seq predicted binding of 36 chromatin factors with a median MCC *>* 0.3. These 36 chromatin factors had a auPR between 0.27 and 0.84 (Table 1). Some of these chromatin factors show high levels of consistent binding among different cell types, which makes predictions easier. The fraction of bins bound to a chromatin factor in at least half of training cell types, however, varies between 0 to 15.75% across all chromatin factors. Even for chromatin factors with a median auPR *>* 0.5 (purple in Figure 4a) the fraction of bins bound in half of training cell types varied from 0.5% in FOXA1 to 10.5% in NRF1. For some DNA-binding proteins, Virtual ChIP-seq fails to predict binding accurately (auPR < 0.3). DNA-binding proteins with low auPR and low MCC include chromatin modifiers such as KAT2B, KDM1A, EZH2 and chromatin binding proteins such as CHD1 and BRD4. Chromatin factors with low prediction accuracy include ATF2, CUX1, E2F1, EP300, FOSL1, FOXM1, JUN, RCOR1, RELA, RXRA, SREBF1, TCF12, TCF7L2, and ZBTB33. For some proteins, such as ATF2, EP300, EZH2, FOXM1, KAT2B, KDM1A, TCF12, and TCF7L2, in at least one validation cell type, most ChIP-seq peaks didn’t overlap with chromatin accessible regions.

#### 2.2.10 Features of true and false predictions

To better understand why the model sometimes predicted incorrectly, we examined predictions of 52 chromatin factors in validation chromosomes (chr5, chr10, chr15, and chr20) in K562. We investigated true positive (TP), false positive (FP), and false negative (FN) predictions. We excluded true negative (TN) predictions because their high numbers mainly reflect imbalanced class prevalence and potential ascertainment bias in the ground truth. Among the three labels, TP genomic bins varied from 0.19% for RELA to 58% for CTCF (Figure 5a). For 24 of these 52 chromatin factors, most incorrect predictions were FN (Figure 5a, left). For the other 28 chromatin factors, most incorrect predictions were FP (Figure 5a, right).

**Figure 5:**
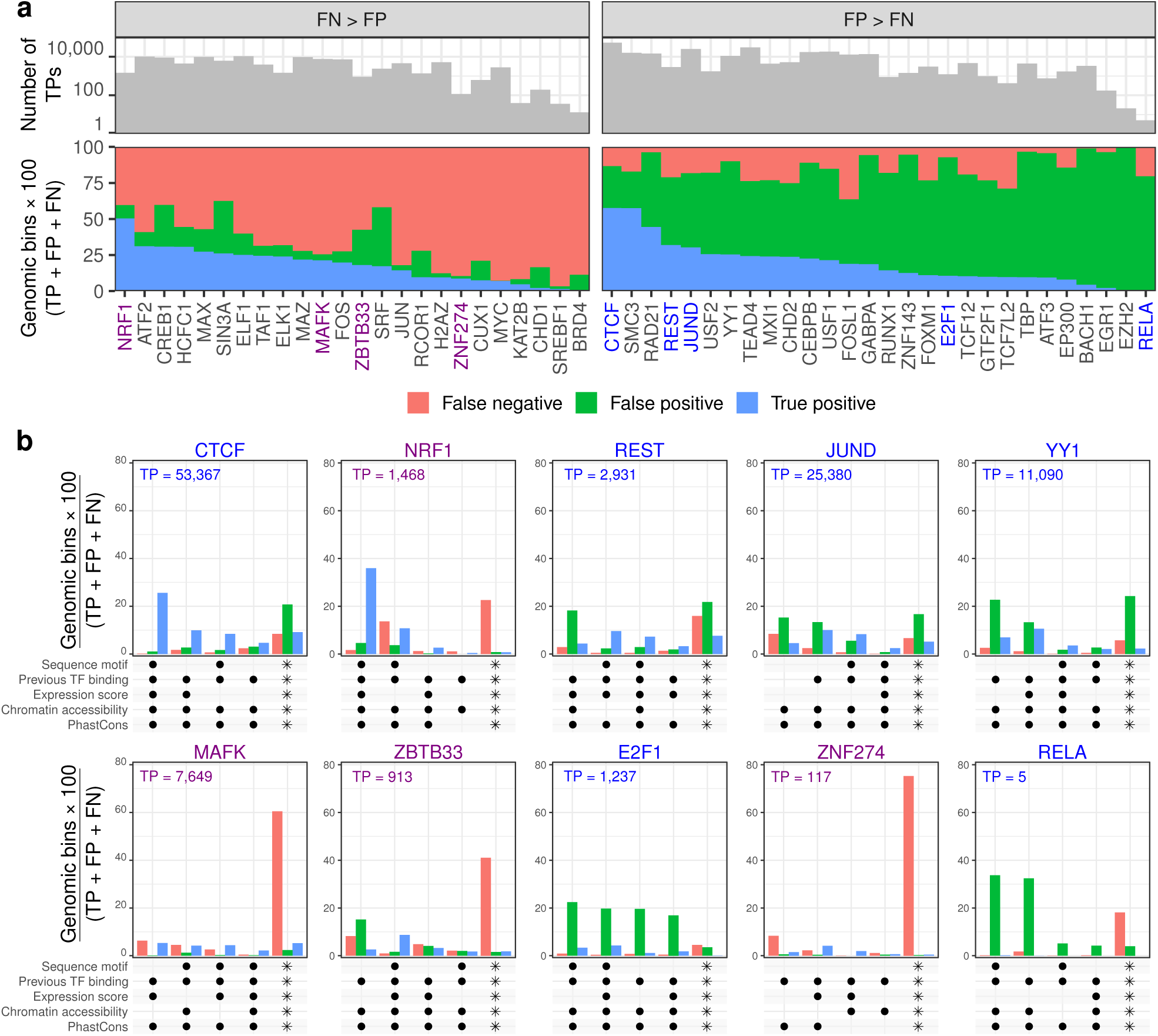
True and false predictions and associated features. Fraction of potential chromatin factor binding sites in K562 categorized as false negative (FN: orange), false positive (FP: green), and true positive (TP: blue). This excludes any sites deemed true negative (TN). **(a)** Stacked bar plot of prediction categorization for the 52 chromatin factors with K562 ChIP-seq data, sorted by the fraction of TP genomic bins. Grey bars show number of TP predictions. (*Left*) 24 chromatin factors where FN fraction exceeded FP fraction. We selected 4 factors to examine in more detail below (orange names). (*Right*) 28 chromatin factors where FP fraction exceeded FN fraction. We selected 6 factors to examine in more detail below (green names). **(b)** UpSet^58^ plot of prediction categorization in 10 factors given the 4 most common combinations of positive values for input features, and all other combinations (asterisks). Black dots indicate the features with positive values in each combination. Number of TPs indicated is in validation chromosomes (chr5, chr10, chr15, and chr20). We took the 10 factors from a wide range of those with best performance (top left) to worst performance, as sorted by ratio of TP to FP+FN.

We investigated presence and absence of predictive features among genomic bins labeled TP, FP, and FN. We defined presence of a feature as a positive value, and absence as a non-positive value. Expression score has values in [−1, 1] when a region had chromatin factor binding in any of the training cell types that have matched RNA-seq data. For expression score, non-positive values include both 0 and negative values. All other input features only have values in [0, 1]. For most chromatin factors, the model performed better when all features were present. This means higher TP, lower FN, and lower FP (Figure 5b).

For CTCF, incorrect predictions represented less than 5% of TPs when all predictive features were present, when only sequence motif was absent, or only the expression score was absent (Figure 5b). Without presence of chromatin accessibility, the model made a higher number of false predictions, but still made some correct predictions.

The model only predicted novel binding sites not present in training cell types when the site matched the TF’s sequence motif (Figure 5b). For NRF1, MAFK, and ZNF274, the model made frequent FN predictions when expression score and sequence motif match were absent. REST, JUND, YY1, and E2F1 have more FP than FN. For these TFs, FP predictions were frequent when expression score and sequence motif match were absent. For ZBTB33, both FP and FN predictions were high when expression score and sequence motif match were both absent.

ZNF274 had only 117 correctly predicted binding sites and RELA had only 5 correctly predicted binding sites in the four validation chromosomes. In both of these cases, the model had low specificity and sensitivity, predicting a much higher number of FNs and FPs than TPs.

#### 2.2.11 The expression score leverages similarity with training cell types to improve predictions

The expression score for a genomic bin is the Spearman correlation between expression of specific genes in a new cell type and a measure of how chromatin factor binding in that genomic bin correlates with expression of those genes among training cell types. For each genomic region, the expression score uses the expression values of a different set of genes to provide a low or high probability for chromatin factor binding in the new cell type.

We investigated whether the expression score serves as a way of encoding the ChIP-seq data of the training cell type with the most similar transcriptome to the new cell type. To do this, we randomly permuted expression scores across the genome. We identified bins that have TP predictions with the original expression score but switch to FN with the permuted score. The correct predictions that require the original expression score usually had ChIP-seq peaks in one or more training cell type. In rare cases, these apparently expression-requiring predictions did not have corresponding binding in any of the training cell types. In these cases, the expression score may have contributed little to original prediction, but a permuted expression score penalized the bin below the prediction threshold.

We investigated the TF JUND in more detail. In JUND, 126 out of 1,155 expression-requiring TP predictions did not exist in any of the training cell types (Figure 6a, blue). Some of these true predictions (117/1,155) existed in only one of the training cell types (Figure 6a, orange). We investigated correlation of the rank of expression of the top 5,000 genes with the highest variance among training cell types and the validation cell type K562 (Figure 6a). The training cell type with the highest correlation was not necessarily the cell type with the highest number of expression-requiring predictions. For example, although the correlation among expression of all of the 5000 genes is highest between HeLa-S3 and K562 (*r* = 0.44), HCT-116 (*r* = *-*0.13) is the source of the highest number of correct expression score specific predictions. This is unsurprising since, for each region’s expression score, we used only a subset of the 5,000 genes in the global calculation here. The other 912 predictions existed in 2 or more training cell types. This implies that, at least for JUND, the expression score did not simply encode ChIP-seq data of a single training cell type with the most similar global transcriptome to the new cell type.

**Figure 6:**
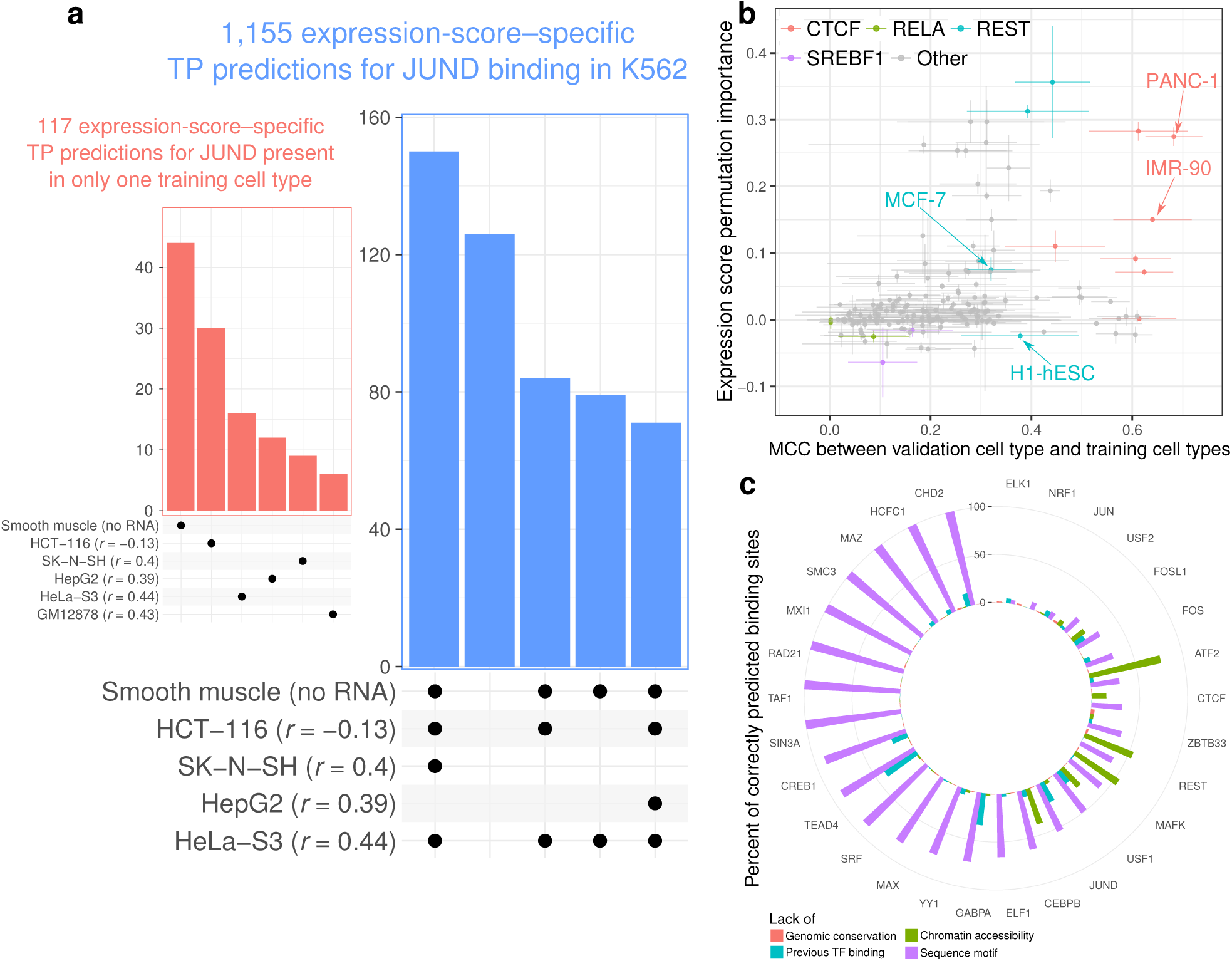
Expression score leverages similarity with training cell types. **(a)** UpSet plots of TP predictions of JUND binding in K562 which did not pass the posterior probability threshold when we permuted the expression score. Each bar represents a combination of training cell types with the binding site (black dots below plot). *r*: genome-wide correlation of rank of expression of the top 5,000 genes with highest variation among a training cell type with rank of expression of the same genes in K562. Smooth muscle lacked matched RNA-seq data. Blue plot: the top 5 combinations with the highest number of TP genomic bins. Orange plot: the TP predictions which were bound to chromatin factor in only one training cell type. **(b)** Scatter plot of expression score permutation importance for 160 pairs of 63 chromatin factors and 6 validation cell types against ChIP-seq peak similarity between that cell type and 1–10 training cell types. Permutation importance is the difference in auPR when permuting expression score. Similarity is measured by MCC of validation cell type ChIP-seq peaks, treating each training cell type in turn as ground truth. Each point indicates median quantities, and error lines indicate median absolute deviation. **(c)** Bar plot of the fraction of binding sites for 29 chromatin factors correctly predicted on K562 validation chromosomes (chr5, 10, 15, and 20) which lacked particular predictive features. These features include genomic conservation (red), chromatin accessibility (green), sequence motif (turquoise), and evidence of chromatin factor binding in another cell type (purple). For chromatin factors with no sequence motif, we deemed every binding site to lack a sequence motif.

We also examined whether the expression score’s effectiveness depends on the similarity of chromatin factor binding among training and validation cell types. Under this hypothesis, we would expect high correlation between the expression score’s contribution to model performance and the similarity of ChIP-seq data between the validation cell type and the training cell types. To examine this hypothesis, we calculated pairwise similarity in ChIP-seq data between the validation cell type and each training cell type. Due to the highly imbalanced class prevalence of ChIP-seq data, we used pairwise MCC as the similarity measure. We also calculated permutation importance^59^, the difference in auPR when permuting the expression score (auPR – auPR_permuted expression score_). Permutation importance indicates a feature’s contribution to a predictive model.

For each validation cell type, we calculated the median MCC of its ChIP-seq data with that of training cell types and median expression score permutation importance among the 4 validation chromosomes (Figure 6a). These two variables correlate in general (Spearman’s *ρ* = 0.41; *p* = *×* 10^−8^). CTCF binding in PANC-1 similarity with training cell types ranges from MCC = 0.38 to MCC = 0.76 (Figure 6b). Only CTCF binding in IMR-90 has a higher similarity to training cell types (MCC *∈* [0.35, 0.79]). The permutation importance of CTCF predictions in PANC-1 is 0.27, while the permutation importance of CTCF predictions in IMR-90 is 0.15. The variation in correlation of similarity to training cell types and permutation importance of the expression score is more evident for REST (Figure 6b). While the median similarity of REST binding with training cell types is 0.32 for MCF-7 and 0.38 for H1-hESC, the permutation importance for REST binding is 0.07 for MCF-7 but –0.02 for H1-hESC.

Using the expression score generally improved performance when validation cell types had similar TF location patterns to training cell types. For example, some validation cell types similar to the training cell types often had high expression score permutation importance (*≥* 0.1) for CTCF (IMR-90, liver, MCF-7, PANC-1) and REST (K562, PANC-1). For RELA and SREBF1, however, all validation cell types had low expression score permutation importance (< 0.1), and low similarity of ChIP-seq data to training cell types (Figure 6b).

#### 2.2.12 Some correct predictions lack known predictive features

Many correctly predicted binding sites in K562 lack important predictive features of chromatin factor binding (Figure 6c). Among 29 chromatin factors with MCC *>* 0.3 in K562, almost all correct predictions are in genomic bins conserved among placental mammals^47^,48. The exceptions include 3.72% of predictions for ZBTB33, 2.11% of predictions for REST, 2.07% of predictions for USF2, 1.49% of predictions for NRF1, 1.47% of predictions for CHD2 and 0.18%–0.89% for other chromatin factors. Many correctly predicted binding sites for ATF2, MAFK, REST, CEBPB, USF1, FOSL1, and CTCF don’t overlap chromatin accessibility peaks. We correctly predicted many binding sites for TEAD4, GABPA, JUND, CREB1, USF1, CHD2, and FOSL1 in regions which had no binding in training cell types. For all these factors except JUND, the nearest upstream or downstream genomic bin of these novel predictions in K562 bound the chromatin factor as well. The nearest training cell type binding site to these correct novel predictions were 50 bp–3.6 Mbp away. The nearest peak in training cell types for these novel predictions was not significantly closer compared to other K562 ChIP-seq peaks (Wilcoxon rank sum test; *p* = 1). In these cases, the multi-layer perceptron learned from other available predictive features. For example, in TEAD4, all novel correctly predicted binding sites in validation chromosomes overlapped chromatin accessibility peaks. These correct predictions also had a mean PhastCons conservation of 0.182, significantly higher than the mean of 0.150 in other genomic bins (Welch t-test; *p* < 2 × 10^−16^).

### 2.3 The choice of input features determines prediction performance

#### 2.3.1 The most important features

To evaluate the importance of each feature in our predictive model, we performed an ablation study on training data. First, we systematically removed features. Second, we fitted the model without these features on some of the training cell types (HeLa-S3, GM12878, HCT-116, LNCaP). Third, we evaluated performance on one held-out training cell type (HepG2; Supplementary Table 12). This ablation study did not use any of the validation cell types which we used for final evaluation of the model.

We called the effect of excluding an input feature substantive only when the average increase or decrease in auPR was at least 0.05. Excluding genomic conservation, sequence motif, HINT, or CREAM did not substantively change performance of the model for most chromatin factors (Figure 7). Excluding chromatin accessibility, publicly available ChIP-seq data, and the expression score decreased performance in most chromatin factors. Excluding expression score substantively decreased median auPR in 13/21 chromatin factors, while excluding publicly available ChIP-seq data substantively decreased auPR in 18/21 chromatin factors.

**Figure 7:**
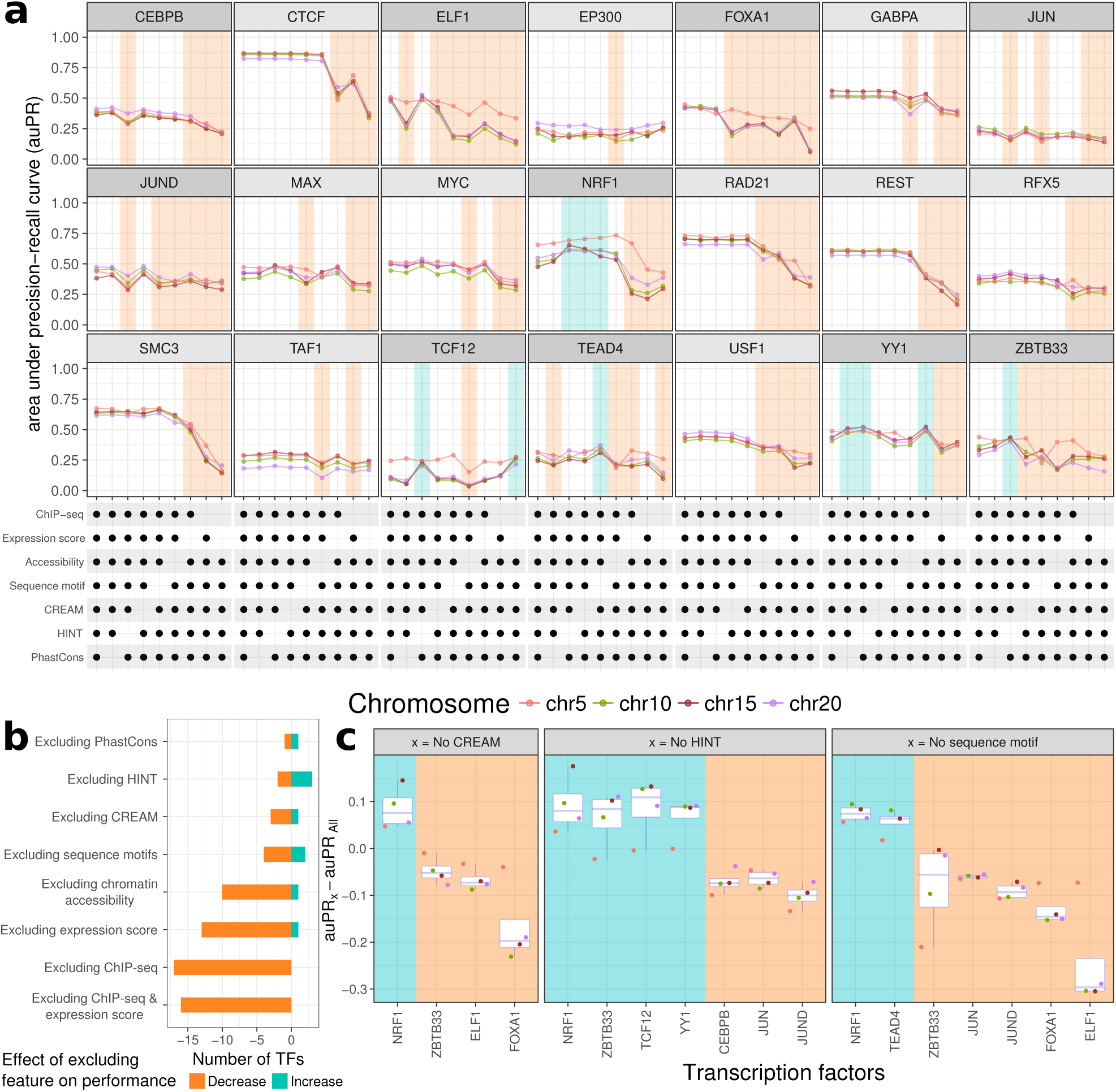
Virtual ChIP-seq’s most important features consist in ChIP-seq data and expression score. **(a)** Area under precision-recall curve (auPR) for predicting a chromatin factor’s binding sites after training on only a subset of input features. We trained on five cell types (HeLa-S3, GM12878, HCT-116, and LNCaP) and predicted on either HepG2. Ablating a feature caused either substantive decrease (orange), substantive increase (turquoise), or no substantive change in auPR. An UpSet^58^-like matrix shows the subset of features used for each column. Dark grey strip above facet: when ablating HINT, CREAM, or sequence motifs substantively changed auPR. **(b)** Double-ended bar plot of the number of chromatin factors with average auPR increase or decrease of at least 0.05 when ablating each feature. Bars show the number of chromatin factors where ablation caused the average auPR to decrease (orange, left) or increase (turquoise, right). **(c)** Change in auPR for those chromatin factors with an average auPR increase or decrease of at least 0.05 when we excluded clusters of regulatory elements (CREAM), footprints (HINT), or sequence motifs. Backgrounds indicate auPR decrease (orange) or increase (turquoise).

#### 2.3.2. Inclusion of some features have opposite effects on prediction of different chromatin factors

Beyond the most important features—chromatin accessibility, ChIP-seq, and expression score— excluding other features rarely substantively decreased prediction performance (Figure 7b–c). When we excluded sequence motifs, auPR decreased substantively for ZBTB33, JUN, JUND, FOXA1, and ELF1. Excluding HINT footprints decreased auPR substantively only for CEBPB, JUN, and JUND. Excluding CREAM clusters of chromatin accessibility peaks decreased auPR substantively only for ZBTB33, ELF1, and FOXA1.

Removing certain input features actually improved prediction for some chromatin factors (Figure 7b–c). Associations that differed between training cell types and validation cell types suggested that these input features generalize poorly. For example, CREAM clusters’ overlap with NRF1 ChIP-seq peaks was not consistent among GM12878 (7.52%), HeLa-S3 (31.8%), and HepG2 (25.78%). This represented a significant variation among these cell types (ANOVA; *p* = 1.9 × 10^−4^).

While most TF footprints (95.96%) overlapped NRF1 peaks, TF footprints constituted only a small fraction of NRF1 peaks (0.73%). NRF1 peaks overlapped a smalls proportion of TF footprints in training cell types GM12878 (1.14%) and HeLa-S3 (0.59%), but significantly greater than the 0.45% overlap in HepG2 (Welch t-test; *p* = 0.007). In HepG2, 7.28% of YY1 peaks overlap TF footprints while in the training cell type GM12878, the overlap is only 1.22% (Welch t-test; *p* = 5 × 10^−5^) and in the other training cell type HCT-116 the overlap is much higher (17.92%; Welch t-test; *p* = 5 × 10^−6^). Overlap of ZBTB33 peaks with TF footprints is much smaller in HepG2 (0.49%) compared to training cell types GM12878 (2.32%) and HCT-116 (5.27%; Welch t-test; *p* = 6 × 10^−4^). Features with varying and cell-specific association with chromatin factor binding complicate convergence of the multi-layer perceptron and may result in overfitting. As a result, the multi-layer perceptron achieved a higher performance on some chromatin factors when we ablated those features.

Association of clusters of regulatory elements and chromatin factor footprints with chromatin factor binding varies among cell types. Using a CREAM feature substantively improved performance in 3/21 chromatin factors and using a HINT feature substantively improved performance in 3/21 chromatin factors (Figure 7b–c). In contrast, including CREAM substantively decreased performance for 1 case and including HINT for 4 cases. When we repeat this experiment by using different training and validation cell types, clusters of regulatory elements and TF footprints result in increase or decrease in performance of different chromatin factors, while they barely result in an increase in auPR above 0.05. Because of the limited upside and apparent downside, we didn’t use these two cell-type–specific features for our final model.

### 2.4 Transcription factors and their targets regulate similar biological pathways

#### 2.4.1 Gene set enrichment analysis of chromatin factor targets

To calculate the expression score, we investigate correlation of chromatin factor binding at each genomic bin with expression of 5,000 genes across the genome (Methods). This brings us to our hypothesis that genes whose expression is perturbed with binding of a chromatin factor regulate the same biological processes as the chromatin factor. To understand biological implications of transcriptome perturbation in response to chromatin factor binding, we measured how frequently each gene’s expression associated with binding of each chromatin factor. We hypothesized that if expression of a gene consistently correlates with binding of a chromatin factor, it is a potential target of that chromatin factor. Similarly, if the expression of a gene negatively correlates with binding of a chromatin factor, cellular machinery upregulated by that chromatin factor might cause net suppression of that gene’s expression.

To identify such genes, for each chromatin factor, we ranked genes by subtracting the number of genomic bins they are positively correlated with from the number of genomic bins they are negatively correlated. We call this difference the *association delta*. For each chromatin factor, we identified the 5,000 genes with the highest variance in expression among cells with matched RNA-seq and ChIP-seq data (Figure 2a). We measured correlation of expression of each of the 5,000 genes with chromatin factor binding at every 100 bp genomic window in 4 chromosomes (chr5, chr10, chr15, and chr20). This approach identified genes that have consistent positive or negative association with chromatin factor binding (Figure 8a). We considered these genes as potential targets of each chromatin factor, and used the Gene Set Enrichment Analysis (GSEA) tool^60^ to identify pathways with significant enrichment in either direction (Figure 8a.) Only the rank of association delta affects these results, and we presumed that there would be little difference in using all chromosomes instead of just 4. The 4-chromosome analysis for JUND had no significant rank difference from an analysis of chromosome 10 alone (Wilcoxon rank sum test *p* = 0.3). We only investigated Gene Ontology (GO) terms annotated to a minimum of 10 and a maximum of 500 out of a total of 17,106 GO-annotated genes.

**Figure 8:**
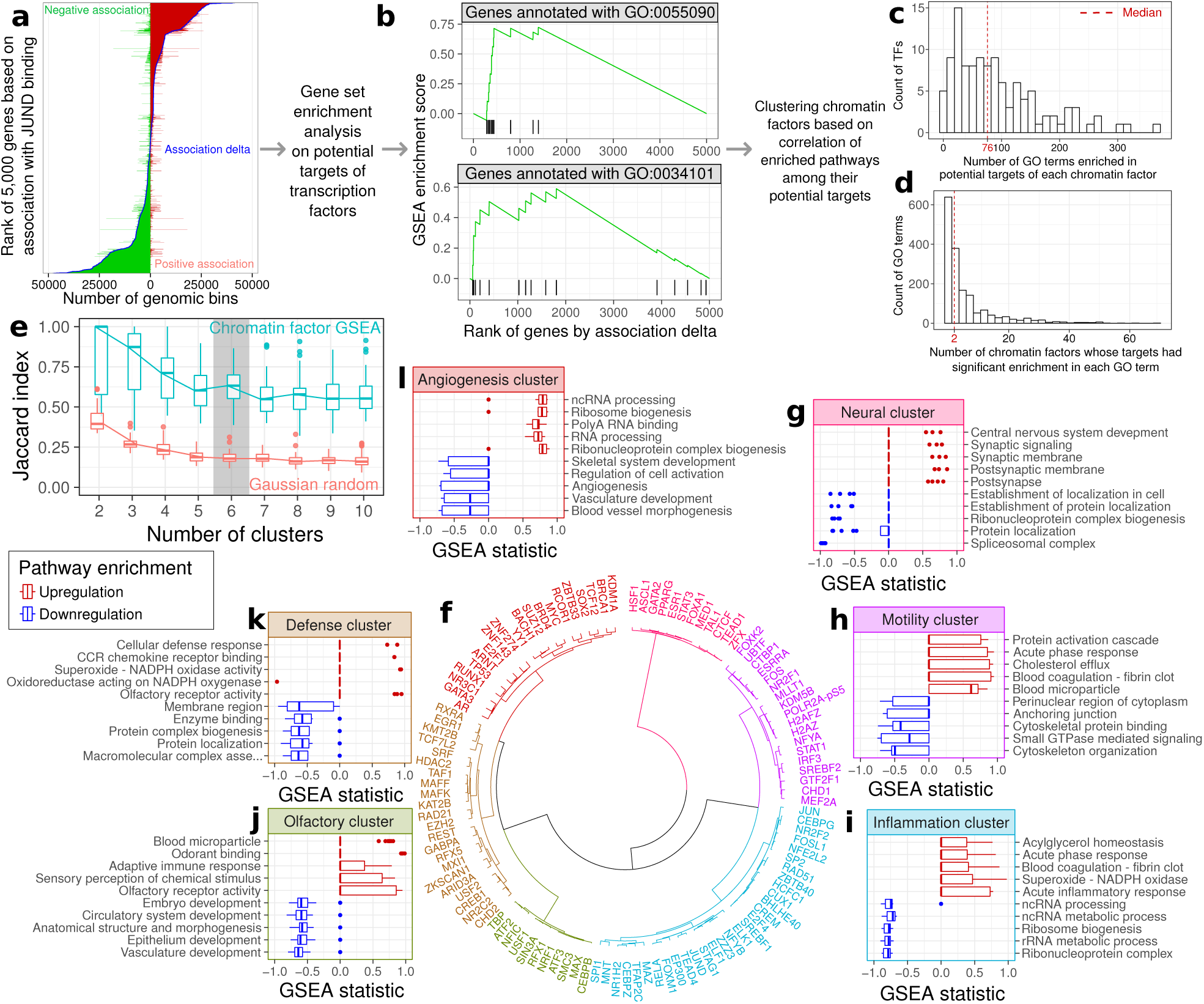
Top biological pathways regulated by potential targets of chromatin factor clusters. Each gene may have both positive and negative correlation with chromatin factor binding at some genomic bins. For each chromatin factor, we ranked 5,000 genes by an association delta that summarizes how many genomic bins associated with binding. The association delta takes the number of bins that positively associated with a gene’s expression and subtracts the number of bins that negatively associated. **(a)** The association ranking process for JUND binding. Double-ended bar plot for each of the 5,000 genes, with positive (red) and negative (green) association. Super-imposed blue curve: association delta for each gene. **(b)** Gene Set Enrichment Analysis (GSEA)^60^ identified pathways with significant enrichment in potential targets of each chromatin factor. Vertical black bars: rank of association delta for genes annotated with each GO term. Green line: GSEA enrichment score. **(c)** Histogram showing how many of 1,681 GO terms are enriched in potential targets of each chromatin factor. **(d)** Histogram showing how many of 63 chromatin factors have potential targets with enrichment in each GO term. **(e)** Boxplot of cluster stability, as measured by Jaccard index, between clusters found in both the subsampled correlation matrix of chromatin factors by GSEA (turquoise) and a subsampled random Gaussian matrix of the same dimensions (red). Grey background: the smallest number of clusters where GSEA matrix cluster stability increased but that of the random Gaussian matrix did not. **(f)** Dendrogram of 6 clusters identified in the correlation matrix. We defined 6 clusters based on correlation of enrichment in 1,681 GO terms. **(g–l)** Boxplots of GSEA statistic for the top 5 pathways enriched in genes positively (red) and negatively (blue) correlated with chromatin factor binding for each cluster.

We identified 1,681 GO terms with significant enrichment (GSEA *p* < 0.001) among potential targets of at least one of the 113 chromatin factors we investigated (Figure 8b). Only 63 of these 113 chromatin factors had matched ChIP-seq and RNA-seq in at least 5 of the training cell types and one of the validation cell types we used for learning from the transcriptome. Each chromatin factor had potential targets with significant enrichment in a mean of 92 terms (median 76; Figure 8c). Each of the 1,681 terms had significant enrichment in potential targets of a mean of 6 chromatin factors (median 2; Figure 8d). Furthermore, 300 of these GO terms had significant enrichment in potential targets of at least 10 chromatin factors.

To identify chromatin factors involved in similar biological processes, we searched for enrichment of any of the 1,681 GO terms in 113 chromatin factors. This analysis relied on the GSEA enrichment score as a normalized test statistic. We examined the pairwise correlation between the vector of enrichment scores for each pair of chromatin factors. These pairwise correlations constitute a symmetric correlation matrix. We hypothesized that chromatin factors with high correlation are involved in similar biological processes.

To identify groups of chromatin factors involved in similar biological processes, we performed hierarchical clustering on the correlation matrix. We sought to identify clusters of chromatin factors, and the best number of clusters between 2 and 10, inclusive. As a control, we generated a correlation matrix of same dimensions from a matrix of random Gaussian values (Methods). For each matrix we repeatedly generated random subsamples and clustered them. For each subsample, we found the set of pairs of chromatin factors with the same cluster membership. For couples of these subsamples, we identified the Jaccard index between these sets as a measure of cluster stability^109^ (Methods). We then compared the increase or decrease in Jaccard indices from each number of clusters to the number of clusters one larger.

The smallest number of clusters with an increase in Jaccard index only for the correlation matrix was 6 (Figure 8e–f). We assigned names to these clusters based on their enriched biological pathways. We then examined the chromatin factors included in those clusters. The Neural cluster (Figure 8g) includes ASCL1^61^, HSF1^66^, GATA2^65^, and PPARγ ^67^. These chromatin factors play important roles in the development of the nervous system and are implicated in neurological disorders^61,65,66,67^. The top 5 GO terms enriched in the potential targets of these chromatin factors are all related to nervous system development and function (Figure 8g). The downregulated pathways of the Motility cluster (Figure 8h) relate to cytoskeletal organization. The included chromatin factors, CTBP1^71^, KDM5B^72^, MEF2A^73^, and STAT1^74^, all play a role in the epithelial-to-mesenchymal transition, which involves re-organization of the cytoskeleton. Similarly, we found that for other clusters, specific upregulated or downregulated pathways of cluster’s targets are also regulated by many of the cluster’s chromatin factors (Figure 8i–l, Table 2).

**Table 2:**
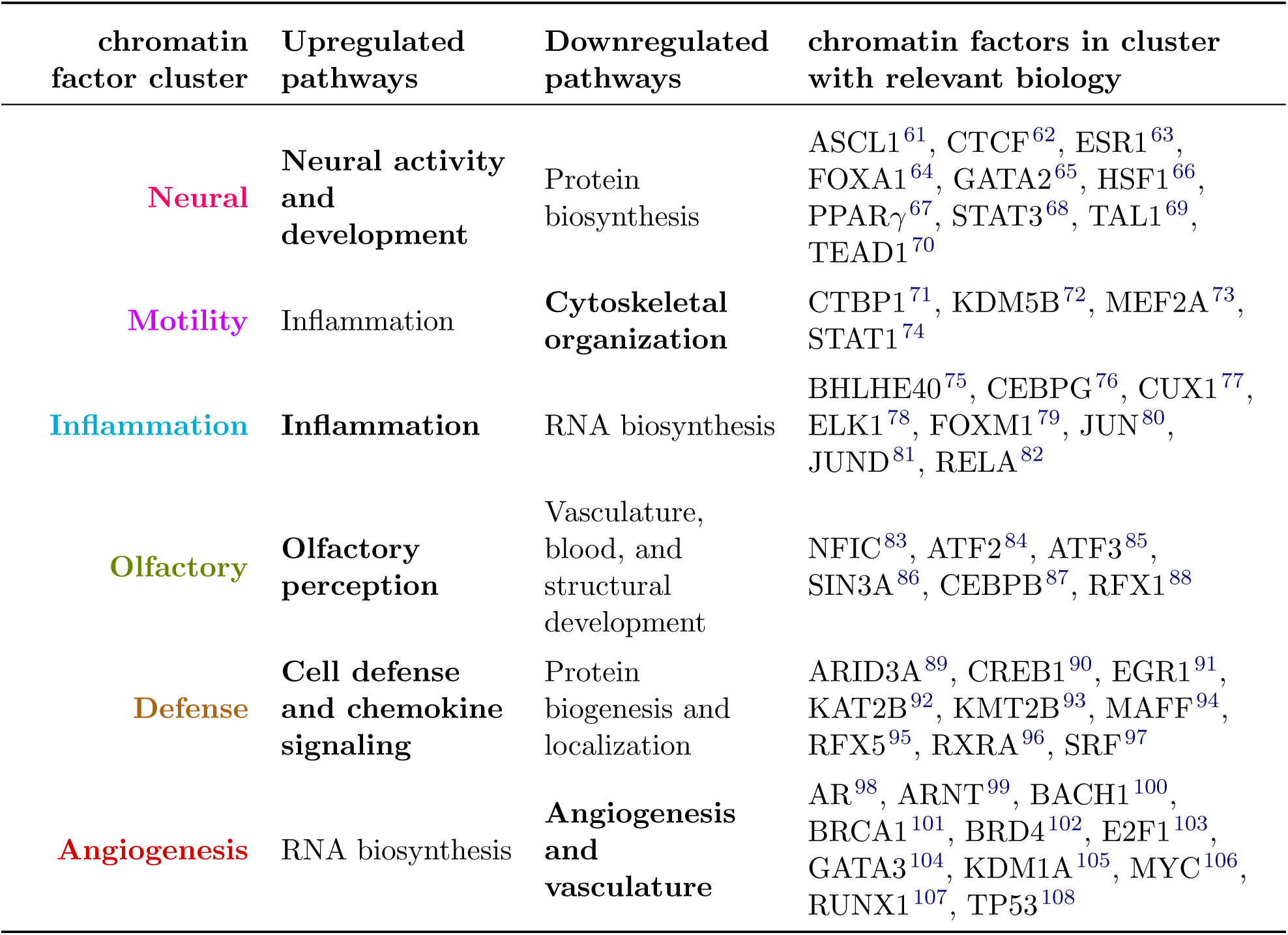
Many chromatin factors within each biological function cluster are involved in the same pathways as their potential target genes. We summarized each cluster of chromatin factors according to top over-represented GO terms in the first 3 columns. Chromatin factors in the 4th column are involved in the same biological mechanism as the bold pathways mentioned in 2nd or 3rd column.

### 2.5 A compendium of chromatin factor binding predictions for 33 tissues and cell types

#### 2.5.1 Predicting chromatin factor binding in Roadmap datasets

The Roadmap Epigenomics Project^35^ performed DNase-seq on 55 and RNA-seq on 39 human tissues and cell types, but not ChIP-seq of any chromatin factor. For 33 of these tissues, they produced matched DNase-seq and RNA-seq data. This makes the Roadmap data an ideal application for Virtual ChIP-seq.

We generated an annotation similar to peak calls by converting the multi-layer perceptron’s posterior probabilities to a presence or absence call. We made this call based on a different cutoff for each chromatin factor. We defined this cutoff as the posterior probability which maximized MCC in H1-hESC. For chromatin factors without ChIP-seq data in H1-hESC, we used the mode of cutoffs from the other different chromatin factors (0.4). We excluded H1-hESC when reporting all performance metrics that depend on this threshold. The number of binding sites we predicted in other validation cell types and Roadmap data is similar to ChIP-seq peaks in other validation cell types (Figure 9a).

**Figure 9:**
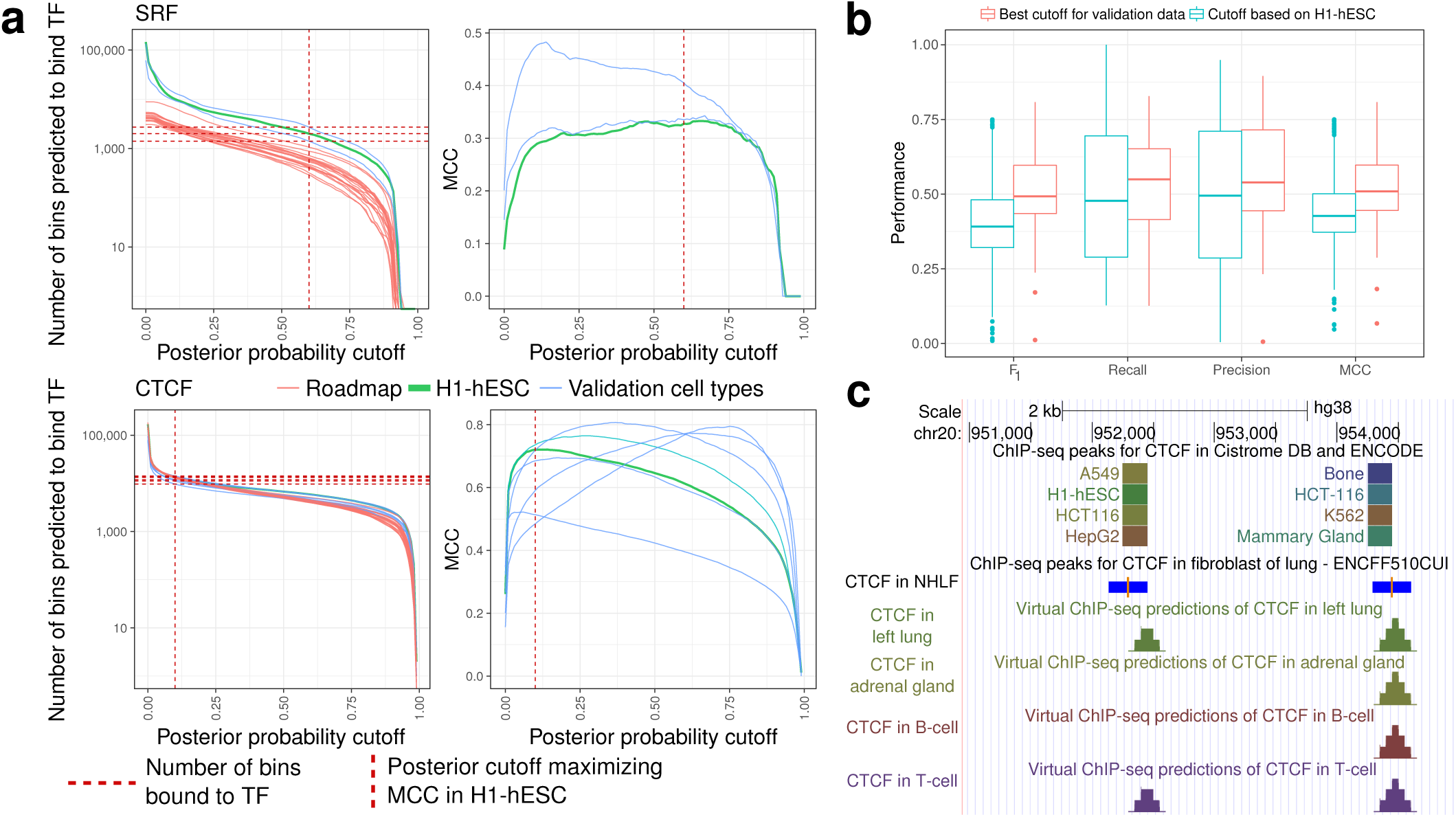
Chromatin factor binding predictions in validation cell types and Roadmap datasets. **(a)** Number of genomic bins that chromatin factor is predicted to bind (left) and MCC (right) as a function of posterior probability cutoff for SRF (top) and CTCF (bottom). This relationship is shown for H1-hESC (turquoise), 2 validation cell types for SRF (blue), and 6 validation cell types for CTCF (blue). Each curve represents predictions for one of the 4 chromosomes (chr5, chr10, chr15, and chr20). Left panels also show how many genomic bins are predicted to bind the chromatin factor in 18 Roadmap datasets (red). Vertical red dashed line: posterior probability cutoff which maximized MCC of the chromatin factor in H1-hESC. Horizontal red dashed lines: number of genomic bins with chromatin factor binding in validation cell types. **(b)** Boxplot of various performance measures when using the best cutoff for each dataset (red) and the optimal cutoff in H1-hESC (turquoise). **(c)** UCSC Genome Browser display of a 4000 bp region on chromosome 20 using the Virtual ChIP-seq track hub (https://virchip.hoffmanlab.org). The track hub has a supertrack for each chromatin factor. Each supertrack contains 34 stracks: one track specifying genomic bins bound by that chromatin factor in Cistrome and ENCODE, and one track for each of the 33 Roadmap cell types with predictions for that chromatin factor. This example shows parts of the track hub related to CTCF, including a track with experimental results in Cistrome DB and ENCODE with 7 out of 144 cell types enabled, and Virtual ChIP-seq predictions in left lung, adrenal gland, B-cell, and T-cell. The height of predictions indicates the number of overlapping genomic bins predicted to bind the chromatin factor, ranging between 0–4. Between are MACS2 narrow peak calls for CTCF in normal human lung fibroblasts (NHLF) from ENCODE (ENCFF510CUI). Blue: peaks; orange: peak summits.

Using the cutoff which maximized MCC in H1-hESC only slightly decreased performance measurements from what one could achieve with the optimal cutoff for each cell type (Figure 9b). For example, the MCC score showed a median decrease of 0.06 and F_1_ score showed a median decrease of 0.1.

As a community resource, we created a public track hub (https://virchip.hoffmanlab.org) with predictions for 33 Roadmap cell types (Figure 9c). This track hub contains predictions for 36 chromatin factors which had a median MCC > 0.3 in validation cell types (Table 1).

## 3 Methods

### 3.1 Data used for prediction

#### 3.1.1 Overlapping genomic bins

To generate the input matrix for training and validation, we used 200 bp genomic bins with sliding 50 bp windows. We excluded any genomic bin which overlaps with ENCODE blacklist regions (https://www.encodeproject.org/files/ENCFF419RSJ/@@download/ENCFF419RSJ.bed.gz). Except where otherwise specified, we used the Genome Reference Consortium GRCh38/hg38 assembly^50^.

#### 3.1.2 Chromatin accessibility

We used Cistrome DB ATAC-seq and DNase-seq narrowPeak files for assessing chromatin accessibility (Supplementary Table 8). We mapped the signal value of peak summits to all the bins overlapping that summit. In rare cases where a genomic bin overlaps more than one summit, we used the signal value of the summit closest to the p terminus of the chromosome When data were available from multiple experiments, we averaged signal values. Because Cistrome DB does not include raw data that one can use for DNase footprinting, we limited the analysis of HINT TF footprinting and CREAM regulatory element clustering to ENCODE DNase-seq experiments on GM12878, HCT-116, HeLa-S3, LNCaP, and HepG2.

#### 3.1.3 Genomic conservation

We used GRCh38 primate and placental mammal 7-way PhastCons genomic conservation^47,48^ scores from the UCSC Genome Browser^110^ (http://hgdownload.cse.ucsc.edu/goldenPath/hg38/phastCons7way). We assigned each bin the mean PhastCons score of the nucleotides within.

#### 3.1.4 Sequence motif score

We used FIMO^36^ (version 4.11.2) to search for motifs from JASPAR 2016^111^ to identify binding sites of each TF that have the sequence motif of that TF. We used the curated, non-redundant JASPAR database of vertebrate sequence motifs to avoid the unnecessary complexity of having similar redundant motifs. To get a liberal set of motif matches, we used a liberal p-value threshold of 0.001 and didn’t adjust for multiple testing. If the motif for the TF didn’t exist in JASPAR, we used other motifs with same initial 3 letters and counted any TF binding site which had overlap with any of those motifs (Supplementary Table 1).

We also used FIMO and JASPAR 2016 to identify the sequence specificity of chromatin accessible regions. For this analysis, we used a false discovery rate threshold of 0.01%. We used any sequence motif matching the initial 3 letters of a TF as a predictive feature of binding for that TF. For many chromatin factors, more than one motif matched this criteria, and we used all as independent features in the model (Supplementary Table 2).

#### 3.1.5 ChIP-seq data

We used Cistrome DB and ENCODE ChIP-seq narrowPeak files. We only used peaks with FDR < 10^−4^. When multiple replicates of the same experiment existed, we only considered peaks that passed the FDR threshold in at least two replicates. We considered bound only those genomic bins overlapping peak summits. We calculated prevalence of bound bins in each chromosome as

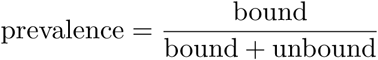

and used it as an auPR baseline^25^.

#### 3.1.6 RNA-seq data

We downloaded an ENCODE expression matrix (https://public-docs.crg.es/rguigo/encode/expressionMatrices/H.sapiens/hg19/2014_10/gencodev19_genes_with_RPKM_and_npIDR_oct2014.txt.gz)^41^ with RNA-seq data for each gene, measured in reads per kilobase per million mapped reads (RPKM). We retrieved similar Cancer Cell Line Encyclopedia (CCLE) RNA-seq data using PharmacoGx^112^. Since these data are processed differently, we limited our analysis to Ensembl gene IDs shared between the two datasets, and ranked gene expression values by cell type. The two datasets have 4 shared cell types: A549, HepG2, K562, and MCF-7. Within each of these cell types, we examined the concordance of RNA-seq data between ENCODE and CCLE after possible transformations. The concordance correlation coefficient^113^ of rank of RPKM (0.827) was higher compared to untransformed RPKM (0.007) or quantile-normalized RPKM (0.006; Welch t-test *p* = 10^−6^). The DREAM Challenge, however, had processed RNA-seq of all cell types uniformly, allowing us to directly use transcripts per million reads (TPM) in analysis of DREAM Challenge datasets.

#### 3.1.7 Expression score

We created an expression matrix for each chromatin factor with matched ChIP-seq and RNA-seq data in *N* ≥ 5 training cell types with the following procedure:

1. We divided the genome into *M* 100 bp non-overlapping genomic bins.
2. We created a non-negative ChIP-seq matrix 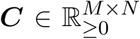 (Figure 2a). We used signal mean among replicate narrowPeak files generated by MACS2^114^ for each of *M* bins and *N* cell types and quantile-normalized this matrix.
3. We row-normalized ***C*** to ***C ′***, scaling the values of each row between 0 and 1.
4. We identified the *G* = 5000 genes with the highest variance among the *N* cell types.
5. We created an expression matrix 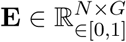 containing the row-normalized rank of expression each of the *G* = 5000 genes in *N* cell types (Figure 2b).
6. For each bin *i* ∈ [1*, M*] and each gene *g* ∈ [1*, G*], we calculated the Pearson correlation coefficient *A*_*i,g*_ between the ChIP-seq data for that bin 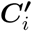, : and the expression ranks for that gene ***E***_:*,j*_ over all cell types. If the Pearson correlation was not significant (*p* > 0.1), we set *A*_*i,g*_ to NA. These coefficients constitute an association matrix ***A*** ∈ (R_*∈*[−1,1]_ *∪* {NA})^*M*×*G*^ (Figure 2c).

We performed power analysis of the Pearson correlation test using the R pwr package^115^.

To predict ChIP-seq binding for a new cell type (Figure 2d), we calculated an expression score for each genomic bin in that cell type. The expression score is Spearman’s *ρ* for expression of the same *G* = 5000 genes in the new cell type with every row of the association matrix ***A***. Each of these rows represents a single genomic bin. An expression score close to 1 indicates that genes with high expression have high values in the association matrix, and genes with low expression genes have low values. An expression score close to −1 indicates that genes with high or low expression have opposite values in the association matrix (Figure 2d).

### 3.2 Training, optimization, and benchmarking

#### 3.2.1 Training and optimization

For the purpose of training and validating the model on Cistrome datasets, we only used chromosomes 5, 10, 15, and 20. These 4 chromosomes constitute 481.78 Mbp (15.6% of the genome). For training only, we excluded any genomic region without chromatin accessibility signal and previous evidence of chromatin factor binding. For validation and reporting performance, we included these regions, using the totality of the 4 chromosomes. We concatenated data from training cell types (A549, GM12878, HepG2, HeLa-S3, HCT-116, BJ, Jurkat, NHEK, Raji, Ishikawa, LNCaP, and T47D; Supplementary Table 3) into the training matrix.

We used Python 2.7.13, Scikit-learn 0.18.1^116^, NumPy 1.11.0, and Pandas 0.19.2 for processing data and training classifiers. We used the default Scikit-learn method^51^ to initialize the multi-layer perceptron’s parameters *β* and coefficients *β*_0_. This uses random values from a uniform distribution. The support of the uniform distribution used depends on properties of the current layer *i* and the next layer *i* + 1. Specifically, the maximum value of the uniform distribution *b* is a function of the number of the hidden units *u*_*i*_ in the current layer, the number of hidden units *u*_*i*+1_ in the next layer, and an activation factor *l* based on the activation function of the current layer. For sigmoid activation, *l* = 2.0 and for other activation functions, *l* = 6.0. For each layer *i*, Scikit-learn sets

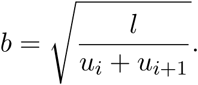

Scikit-learn samples each parameter *β*_*i*_ and each coefficient *β*_0*,i*_ from the uniform distribution 𝒰 (−*b, b*).

We optimized hyperparameters of the multi-layer perceptron^51^ using grid search and 4-fold cross validation. We used minibatch training with 200 genomic bins in each minibatch. We searched for several options to optimize the activation function (Figure 3g), number of hidden units per hidden layer (Figure 3h), number of hidden layers (Figure 3i), and *L*_2_ regularization penalty (Figure 3j). In each round of 4-fold cross-validation, we trained on data of 3 chromosomes, and assessed best MCC on the remaining chromosome. We selected the set of hyperparameters yielding highest average MCC after 4-fold cross validation.

#### 3.2.2 Benchmarking

We used the R precrec package^117^ to calculate auPR and auROC. Precision-recall curves better assess a binary classifier’s performance on imbalanced test data than ROC^25,54^.

#### 3.2.3 DREAM Challenge comparison

For comparison to DREAM results, we also trained and validated the Virtual ChIP-seq model on GRCh37 DREAM Challenge data. For training the model on DREAM Challenge datasets, we used the data of chr5, chr10, chr15, and chr20 of training cell types. We evaluated performance against the union of the DREAM validation chromosomes (chr1, chr8, and chr21) in validation cell types. For CTCF, we trained on all cell types except MCF-7, PC-3, and iPSC which we used for validation. For MAX, we used all cell types except liver and K562 for training. For GABPA, REST, and JUND, we used all cell types except liver for training. We compared these metrics to those of DREAM Challenge participants in the final round of cross–cell-type competition.

### 3.3 Clustering chromatin factors based on enrichment of their potential targets in GO terms

To identify groups of chromatin factors involved in similar biological processes, we performed hierarchical clustering on the correlation matrix. We sought to identify clusters of chromatin factors, and the best number of clusters between 2 and 10, inclusive. For use in this process, we created a Gaussian random matrix of 1,681 rows and 113 columns as a control, and calculated its correlation matrix. Then, we compared cluster stability between the original correlation matrix and the control for each potential number of clusters. To do this, we subsampled 75% of each correlation matrix rows twice without replacement. Then, we clustered chromatin factors in each matrix into the specified number clusters. For both of these clusterings, we constructed the set of every pair of chromatin factors present in the same cluster. We then calculated the Jaccard index between the first clustering’s constructed set and that of the second^109^. We repeated this subsampling and clustering process 50 times for each number of clusters. We picked the smallest number of clusters which had an increase in Jaccard index compared to the number of clusters one smaller only in the chromatin factor correlation matrix.

### 3.4 Chromatin factor prediction on Roadmap data

We downloaded Roadmap DNase-seq and RNA-seq data aligned to GRCh38 from the ENCODE DCC^35^. For each DNase-seq narrowPeak file with matched RNA-seq, we predicted binding of 36 chromatin factors with MCC *>* 0.3 in validation cell types (Table 1, Supplementary Table 6, https://virchip.hoffmanlab.org).

### 3.5 Colors

For plots with three categories, we used a color palette optimized for viewers with deuteranopia (http://mkweb.bcgsc.ca/colorblind) and chose colors also distinguishable by those with protanopia and tritanopia.

For other plots, we either used the default ggplot2^118^ color palette or manually-adjusted ColorBrewer^119^ palettes.

## 4 Discussion

Performing functional genomics assays to assess binding of all chromatin factors may never be possible in patient tissues. Nevertheless, computational prediction of chromatin factor binding based on sequence specificity has identified the role of many chromatin factors in various diseases^1^. Scanning the genome for occurrences of each sequence motif, results in a range of 200–2000 predictions/Mbp. In some cases, this is 1,000 times more frequent than experimental data from ChIP-seq peaks. Similar observations led to a *futility conjecture* that almost all TF binding sites predicted in this way will have no functional role^120^.

Nevertheless, there is more to TF binding than sequence preference. Most chromatin factors don’t have any sequence preference^9^ (Figure 1), and indirect TF binding through complexes of chromatin-binding proteins complicates predictions based solely on sequence specificity. In addition to the high number of false positive motif occurrences, many ChIP-seq peaks lack the TF’s sequence motif. Therefore, relying on sequence specificity alone not only generates too many false positives, but also many false negatives. We call this latter observation the *dual futility conjecture*, although it differs in degree from the original. Adding additional data about cellular state allows us to move beyond both conjectures.

We can assess chromatin factor binding through ChIP-seq or its more precise variations ChIP-nexus^12^ or ChIP-exo^11^. These experiments may still not properly reflect *in vivo* chromatin factor binding due to technical difficulties such as non-specific or low affinity antibodies^121^ or false detection of unrelated factors in hyper-ChIPable regions^122^. Using publicly available ChIP-seq data produced with different protocols and reagents, complicates prediction of chromatin factors more sensitive to experimental conditions^56^. Variations among training and validation cell types in our datasets, overfitted the multi-layer perceptron to certain input features of some chromatin factors. More robust approaches in assessment of chromatin factor binding—such as CRISPR epitope tagging ChIP-seq (CETCh-seq)^123^, which doesn’t rely on specific antibodies—may provide less noisy reference data for learning and prediction of chromatin factor binding.

Virtual ChIP-seq predicted binding of 36 chromatin factors in new cell types, using from the new cell types only chromatin accessibility and transcriptome data. By learning from direct evidence of chromatin factor binding and the association of the transcriptome with chromatin factor binding at each genomic region, most use of sequence motif scores becomes redundant. As more ChIP-seq data in diverse cell types and tissues becomes available, our approach allows predicting binding of more chromatin factors with high accuracy. This is true even in the case of factors that are not sequence-specific. Although Virtual ChIP-seq uses direct evidence of chromatin factor binding at each genomic region as one of the input features, it is able to correctly predict new peaks which don’t exist in training cell types. For 39 of 41 sequence specific chromatin factors, Virtual ChIP-seq correctly predicted chromatin factor binding in regions without any match to sequence motifs.

Virtual ChIP-seq’s performance varies over different chromatin factors, and for each chromatin factor it varies over different genomic regions. When all predictive features had positive values, for example, model performance exceeded conditions where some features were absent. Post-translational modifications to chromatin factors, which none of our input features assess, might explain the varying performance of our model. For example, Virtual ChIP-seq predicts both MYC (MCC = 0.03) and RUNX1 (MCC = 0.27) poorly, and post-translational modifications are known to influence their activity^124,125^. But post-translational modifications also influence the activity of well-predicted chromatin factors CTCF (MCC = 0.68) and SMC3 (MCC = 0.73)^126,127^. Incorporating post-translational modification information remains a future challenge for building more accurate models of chromatin factor binding.

The DREAM Challenge datasets provide data for training and validating machine learning models for predicting binding of 31 chromatin factors. Our datasets, using a combination of Cistrome DB and ENCODE, allow training and validating models for predicting binding in a more extensive 63 chromatin factors. Our provided predictions of binding of 36 high-confidence chromatin factors in 33 different Roadmap tissue types will allow the research community to better investigate epigenomics of disease affecting those tissues (https://virchip.hoffmanlab.org/). In addition to providing our predictions as a resource for use by biologists, we also provide the processed datasets we use as a resource for machine learning researchers. This should accelerate the development of future methods by many groups.

## Acknowledgments

We thank X. Shirley Liu (ORCID: 0000-0003-4736-7339) for providing the Cistrome DB narrow-Peak files. We thank the Roadmap Epigenomics Mapping Consortium and the ENCODE Project Consortium for generating the datasets which enabled this work. We thank Sage Bionetworks-DREAM and the ENCODE-DREAM Challenge organizers for providing data and results before publication. We thank Carl Virtanen and Zhibin Lu (University Health Network High Performance Computing Centre and Bioinformatics Core) for technical assistance. We thank Anshul Kundaje (ORCID: 0000-0003-3084-2287), Nicolae R. Zabet (ORCID: 0000-0001-9964-6271), Patrick Martin (ORCID: 0000-0002-4093-8277) and those at Banff International Research Station Workshop on “The Role of Genomics and Metagenomics in Human Health: Recent Developments in Statistical and Computational Methods” for comments on this manuscript. This work was supported by the Canadian Cancer Society (703827 to M.M.H.), the Ontario Ministry of Training, Colleges and Universities (Ontario Graduate Scholarship to M.K.), and the University of Toronto Faculty of Medicine Frank Fletcher Memorial Fund (M.K.).

